# Variation in phosphorus and sulfur content shapes the genetic architecture and phenotypic associations within wheat grain ionome

**DOI:** 10.1101/580423

**Authors:** Andrii Fatiukha, Valentina Klymiuk, Zvi Peleg, Yehoshua Saranga, Ismail Cakmak, Tamar Krugman, Abraham B. Korol, Tzion Fahima

## Abstract

Dissection of the genetic basis of ionome is crucial for the understanding of the physiological and biochemical processes underlying mineral accumulation in seeds, as well as for efficient crop breeding. Most of the elements essential for plants are metals stored in seeds as chelate complexes with phytic acid or sulfur-containing compounds. We assume that the involvement of phosphorus and sulfur in metal chelation is the reason for strong phenotypic associations within ionome. Thus, we adjusted element concentrations for the effect of variation in phosphorus and sulfur seed content. The genetic architecture of wheat grain ionome was characterize by QTL analysis using a cross between durum and wild emmer wheat. Adjustment for variation in P and S drastically changed phenotypic associations within ionome and considerably improved QTL detection power and accuracy, resulting in identification of 105 QTLs and 437 QTL effects for 11 elements. A search for candidate genes revealed some strong functional associations of genes involved in transport and metabolism of ions and elements. Thus, we have shown that accounting for variation in P and S is crucial for understanding of the physiological and genetic regulation of mineral composition of wheat grain ionome and can be implemented for other plants.

## Introduction

Metabolism of mineral nutrients plays a crucial role in regulation of most of the biochemical and physiological processes in plants. The ionome is defined as the composition of mineral nutrient and trace elements in a single cell, tissue, organ, or the whole plant (Salt *et al.*, 2008). Seeds comprise a major part of the plant reproductive structure and represent developmental end point that can recapitulate genetic and environmental effects that influenced the ionome of a certain organism during its life cycle (Baxter *et al.*, 2014). Furthermore, mineral nutrients of food crop seeds are essential part in human nutrition. For example, deficiency in zinc and iron in human diet was defined as ‘hidden hunger’ and affects about 2 billion people worldwide, mainly in developing world (Gregory *et al.*, 2017; Cakmak & Kutman, 2018).

The list of mineral elements essential for plants is comprised of macroelements, such as nitrogen (N), phosphorus (P), potassium (K), sulfur (S), calcium (Ca), magnesium (Mg), and microelements, such as iron (Fe), manganese (Mn), copper (Cu), zinc (Zn), molybdenum (Mo), boron (B), chloride (Cl) and nickel (Ni) (Kirkby, 2012). Plants uptake and accumulate also a number of non-essential elements, such as aluminum (Al), arsenic (As), cadmium (Cd), cobalt (Co), selenium (Se), strontium (Sr), and rubidium (Rb) (Kirkby, 2012), which can be toxic at high concentrations. Ionome homeostasis is regulated by genetic and physiological mechanisms (Baxter, 2009), which are strongly dependent on the environmental conditions (Asaro *et al.*, 2016), such as soil mineral composition. Understanding of the genetic regulation of the seed elemental network is a crucial target for basic plant biochemistry and physiology studies, as well as for crop breeding.

Positive associations within cereal grain ionomes were reported recently (Shakoor *et al.*, 2016; Pandey *et al.*, 2016), calling for identification of the underlying factors. One of the explanations of this phenomenon at the genetic level is pleiotropic effect of specific genes (Tester, 1990); another one is “dilution effect” (Davis, 2009; Shewry *et al.*, 2016) when the increase in accumulation of carbohydrates dilutes the concentration of minerals in the seeds. From biochemical point of view, positive associations can be explained by similarity of the chemical nature of the elements (Pauli *et al.*, 2018).

Homeostasis of the critical elements for plant growth, such as N, P and S, affects all plant developmental stages (Amtmann & Armengaud, 2009). However, towards the end of the plant’s life cycle, organic compounds and minerals are remobilized and stored in the seeds to ensure their successful germination (Galili, 1997; Raboy, 1997). N, P and S are playing crucial role in seed metal accumulation, since they compose the chelate complexes that are storing the majority of the essential elements (Rauser, 1999). For example, 80% of P is stored in seeds as phytic acid known to be a major metal chelator (Lott *et al.*, 2000). N is known as a major element for building storage proteins in cereal grains (Muench & Okita, 1997), but also play an important role in metal chelation (Rauser, 1999). Furthermore, S is serving as a major element of S-rich storage proteins (Shewry *et al.*, 2001), and was found in many compounds that were shown to be tightly linked to trace element homeostasis (Na & Salt, 2011), such as phytochelatins and metallothioneins (Cobbett & Goldsbrough, 2002).

Biofortification is the strategy used for improvement of nutritionalquality and mineral content in food crops through genetic enhancement (Neeraja *et al.*, 2017; Bouis & Saltzman, 2017). Biofortification is mostly focused on the increase of grain protein (N), Fe and Zn contents, by using genetic and agronomic approaches (Cakmak *et al.*, 2010; Tiwari *et al.*, 2016), with less attention given to the complex relationships between these target element and other elements (White & Broadley, 2009). The ionomics approach can improve the efficiency of biofortification, by providing a better understanding of the relationships between elements, and avoiding the accumulation of antinutrient components (Thompson, 1993) and simultaneously minimizing the increase of concentration of non-essential elements (Schroeder *et al.*, 2013).

Association analysis is increasingly used for identification of chromosomal regions affecting mineral accumulations (Alomari *et al.*, 2017; Kumar *et al.*, 2018), and ionomic traits (Shakoor *et al.*, 2016; Ziegler *et al.*, 2018). Yet, quantitative trait loci (QTL) analysis, based on segregating mapping populations, remains an important approach for genetic dissection of elemental accumulation (Gu *et al.*, 2015; Velu *et al.*, 2017; Huang *et al.*, 2017). The recent development of wheat high-throughput SNP genotyping assays (Wang *et al.*, 2014) and the assembly of reference genomes for wild emmer wheat (*Triticum turgidum ssp. dicoccoides*, WEW) (Avni *et al.*, 2017) and hexaploid bread wheat (*T. aestivum*) (Appels *et al.*, 2018) allow linking between QTL analysis and annotated genome assemblies for the identification of candidate genes (CG). These advanced resources serve as the basis for fast fine mapping and cloning of target genes, as well as for understanding of their involvement in physiological and biochemical processes.

WEW is the progenitor of cultivated wheat and its germplasm is an important source for genetic variation of agronomic traits (Huang *et al.*, 2016; Klymiuk *et al.*, 2019), including nutrient and mineral quality (Chatzav *et al.*, 2010). Tetraploid durum wheat (*T. turgidum ssp. durum*, 2*n* = 4*x* = 28, BBAA) is employed as a bridge (Klymiuk *et al.*, 2019) for transferring of exotic alleles from WEW into hexaploid bread wheat (2*n*=6*x*=42, BBAADD). We previously evaluated WEW genotypes for grain mineral concentrations (Gomez-Becerra *et al.*, 2010a) and identified QTLs affecting grain mineral content in a RIL population derived from a cross between durum wheat Langdon (hereafter LDN) and WEW accession G18-16 (L×G) using an old marker systems of SSRs and DArTs (Peleg *et al.*, 2009a). Recent SNP genotyping of L×G RILs allowed us to improve the quality of the genetic map with over four-fold higher amount of skeletal (framework) markers (1,369 vs. 307 in previous map) and five-fold smaller average interval length between adjacent markers (1.3 cM vs.7.5 cM) (Fatiukha *et al.*, 2019).

The main goal of this study was to dissect the genetic basis of wheat grain ionome, while accounting for interdependent factors that can hide possible genetic variation and shape the elemental composition in grains. Comprehensive phenotyping of 11 grain elements and 17 complex traits related to yield, morphology, phenology and physiology was obtained under contrasting water regimes for L×G RIL population and allowed us to uncover three factors that are strongly associated with all elements: i) yield, ii) P, and iii) S concentrations. We assume that the involvement of P and S in metal chelation is the reason for their associations within ionome, thus, we propose to use them as estimators of chelators. Mapping of the traits adjusted for variation in P and S resulted in a considerable improvement of QTL detection power and accuracy. The obtained results showed that this adjustment is crucial for understanding of the physiological and genetic regulation of mineral composition of wheat grain ionome. We further used the whole genome assembly of WEW (Avni et al., 2017) for identification of promising candidate genes (CGs) that are associated with the ionome traits.

## Materials and Methods

### Plant material and growth conditions

The RIL L×G population derived from a cross between durum wheat (*T. durum*, cv. LDN) and WEW (accession G18-16) has been described previously (Peleg *et al.*, 2009b; Fatiukha *et al.*, 2019)). The population (150 F_6_ RILs) was grown under three environments across two growing winter seasons (2004–2005 and 2006–2007) in an insect-proof screen-house at the experimental farm of the Hebrew University of Jerusalem in Rehovot, Israel. The field soil was brown-red degrading sandy loam, composed of 76% sand, 8% silt and 16% clay. Two irrigation regimes were applied using drip water system in a first year of experiment, well-watered (WW05, 750 mm) and water-limited (WL05, 350 mm), and only well-watered regime (WW07, 720mm) was applied in a second year of experiment (for further details see Peleg *et al*., 2009).

### SNP genotyping and high-density genetic map

Single nucleotide polymorphism (SNP) genotyping of 150 F_7_ lines was performed using the Illumina Infinium 15K Wheat platform, developed by TraitGenetics, Gatersleben, Germany (Muqaddasi *et al.*, 2017), consisting of 12,905 SNPs selected from the wheat 90K array (Wang *et al.*, 2014). The genetic map contained 4,015 SNPs that represent 1,369 unique loci (skeleton markers) and covered 1835.7 cM (for more details see Fatiukha *et al.*, 2019).

### Phenotypic traits

Seed samples were digested in a closed microwave system and phenotyped for the elemental concentrations of Al, Ca, Cu, Fe, K, Mg, Mn, P, S, and Zn using inductively coupled plasma-optical emission spectroscopy (ICP-OES; Vista-Pro Axial; Varian Pty Ltd, Australia). Certified values of reference grain samples, received from the National Institute of Standards and Technology (NIST; Gaithersburg, MD, USA), were used for validation of the measurements. Concentrations of N in the samples were determined using C/N analyzer (TruSpec CN, Leco Co., USA), while grain protein concentration (GPC) was calculated by multiplying N percentage with conversion factor of 5.83 (Merrill & Watt, 1973). Phenotypes of 17 complex traits: grain yield (GY), thousand kernel weight (TKW), kernel number per spike (KNSP), harvest index (HI), spike dry matter (SpDM), total dry matter (TotDM), carbon isotope ratio (δ13C), osmotic potential (OP), chlorophyll content (Chl), flag leaf rolling (LR), culm length (CL), days from planting to heading (DP-H), days from heading to maturity (DH-M), vegetative dry matter (VegDM), spike length (SpL), flag leaf length (FLL), and flag leaf width (FLW) were collected in the first year of the experiment, as described in Peleg *et al.*, (2009b) and Fatiukha *et al.*, (2019).

Adjusted trait values of 11 elements were obtained by calculating the residuals of a linear regression between the means of the corresponding initial trait values and mean values of ‘adjusting’ factors (subsequently GY, P and S), in order to reduce the biases caused by variations in productivity and the structural elements P and S. GPC adjusted for variation in GY was defined, according to Oury & Godin (2007), as grain protein deviation (GPD). QTL analysis was conducted for four sets of phenotypic traits: the set of initial traits and three sets of adjusted traits.

### Statistical analysis of phenotypic data

The BioVinci software (BioTuring, San Diego, CA, USA) was used for statistical analyses, including correlation, regression, principal component (PCA) and analyses of variance (ANOVA). Phenotypic values of initial and adjusted traits were tested for normal distribution. Correlation network analysis was conducted with the Software JASP 0.9 (JASP Team).

### QTL analysis

QTL analysis was performed with MultiQTL software package (http://www.multiqtl.com) using the general interval mapping (IM) procedure. Single-QTL and two-linked-QTL models were used for detection of genetic linkage for each trait in each environment, separately (Korol *et al.*, 2009). Multi-environment analysis (MEA) was conducted by joint analysis of trait values scored in three environments (WL05, WW05 and WW07). Multiple interval mapping (MIM) was used for reducing the residual variation for each QTL under consideration, by taking into account QTLs that reside on other chromosomes (Kao *et al.*, 1999). The significance of the detected QTL effects was tested with 3000 permutation runs. Significant models were further analyzed by 3000 bootstrap runs to estimate the standard deviations of the chromosomal position and QTL effect. In the case of overlapping QTL effects, the detected QTL was designated as multi-trait QTL, while the co-location of effects on the initial and adjusted traits was considered as a single-trait QTL.

### Candidate gene analysis

The annotated gene models of ‘Zavitan’ genome assembly (Avni *et al.*, 2017) were used to generate a list of genes residing within each QTL interval (1.5 LOD support interval of QTL effect with the highest LOD score), based on the physical positions of SNP markers that were previously assigned by Fatiukha *et al.*, (2019). Potential candidate genes were selected from this list using functional annotations associated with ‘transport’ and ‘metabolism’ of ions and elements.

## Results

### Phenotypic variation in GPC and element concentrations

All 11 initial traits exhibited a wide range and transgressive segregation for the RIL population under three environments (WL05, WW05 and WW07) over 2 years (Figure 1, Figure S1 and Table S1). Most of the traits approximately fit normal distribution under all environments, except for Al under WW07 (Figure S1). ANOVA showed highly significant effects (P≤0.001) of environments and years for all traits (Table S1). Traits under WW05 showed reduction in concentrations of most of the elements, except for Mg and Mn, compared to WL05. A comparison between the two well-watered environments (WW05 and WW07) showed lower concentrations for most of the elements, except for Al, Cu and Mg, across RILs under WW07 (Figure 1). Interestingly, the wild parent (G18-16) showed higher or similar concentrations of elements compared to the cultivated parent in well-watered environments for most of the traits, and lower for some of the elements under water-limited conditions (Table S1). Genotype effect was highly significant (P≤0.001) for most of the traits across environments, except for Al under WL05 and WW05 (0.01<P<0.05), and non-significant for Mn under WL05 (Table S1). Traits adjusted to variation in P and S showed similar ranges, although the traits adjusted to variation in GY showed more narrow ranges compared to the traits adjusted to variation in P and S (Table S2).

**Fig. 1.**
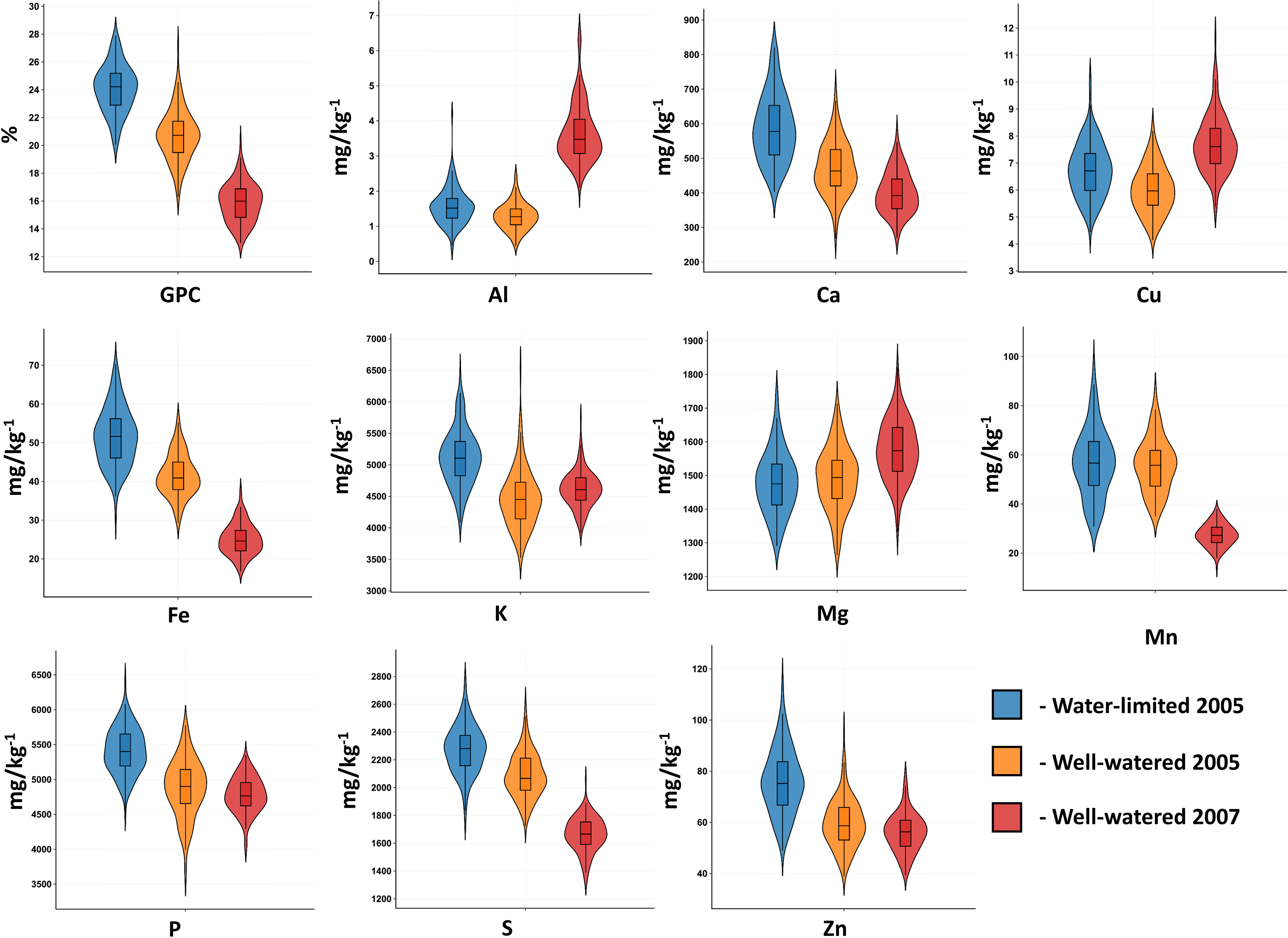
Distribution of 11 wheat grain ionome traits in G×L RIL population. The violin plots are showing the distribution of protein and elemental concentrations in wheat grains measured across three environments WL05, WW05 and WW07

### Phenotypic associations within wheat grain ionome

Correlation analysis within ionome showed strong positive associations between most of the initial traits under all environments (see correlation networks in Figure 2, Table S3-S5). Three traits (Mg, P, and Zn) were positively correlated with all of the other traits, and GPC was not associated only with Al under WL05. In WW05 only Mg displayed significant positive correlations with all other traits, while Al and Mn showed lower associations with other traits in WW05 compared to WL05. The correlation analysis revealed two main differences in associations under WW07 compared to WW05: (i) only three significant correlations between K and other elements, and (ii) more strong associations of Mn with other traits. Although significant effects of irrigation and years on traits were obtained by ANOVA, the associations between elements showed common patterns across all environments.

**Fig. 2.**
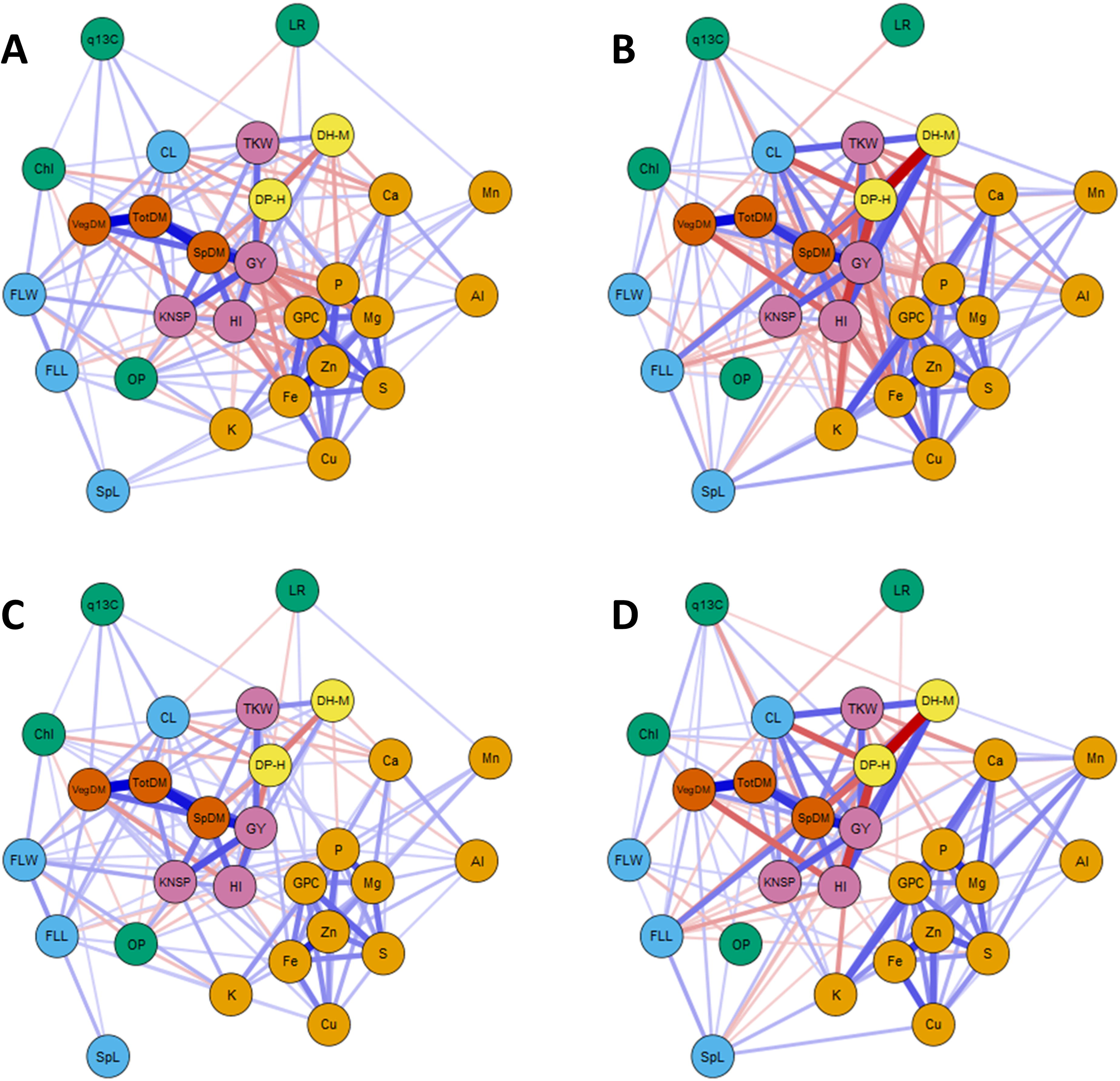
Correlation networks of initial and adjusted to variation in GY ionome traits with 17 complex traits. Networks were obtained for the initial traits under WL05 (A) and WW05 (B) and for the adjusted traits under WL05 (C) and WW05 (D).

We have applied adjustment of GY variation to avoid such potential biases associated with “dilution effect” (Davis, 2009). Unexpectedly, adjustment for variation in GY did not change the associations within the ionome (Figure 3, Table S6-S7). However, adjustment for variations in P and S has resulted in a strong reduction in the associations within the ionome (Figure 3, Table S8-S10), supporting our idea that these two elements play strong unifying effect on other elements and can be used for estimation of the amount of storage chelators in grains. The adjustment for variation in P caused a stronger relaxation in correlations than that of S, although some of the associations continued to be tight, e.g., for Fe-Zn-Cu-GPC. In addition, this adjustment uncovered negative associations between K concentrations and some other elements. In addition, both adjustments showed some environment- and trait-specific associations (Figure 3).

**Fig. 3.**
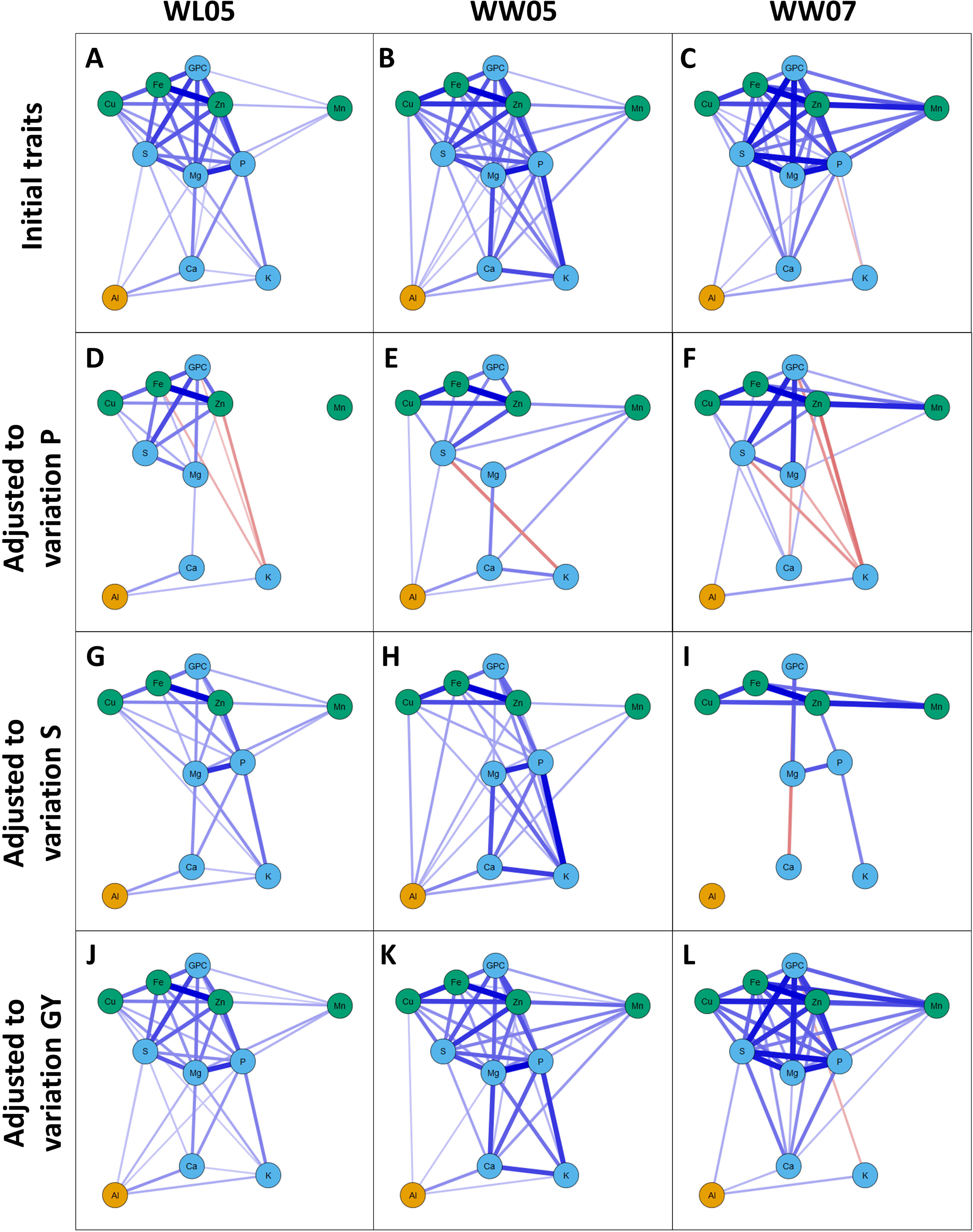
Correlation networks within wheat grain ionome for the initial and the adjusted traits. The networks were obtained for the initial traits under WL05 (A), WW05 (B) and WW07 (C); for the traits adjusted to variation in P under WL05 (D), WW05 (E) and WW07 (F); for the traits adjusted to variation in S under WL05 (G), WW05 (H) and WW07 (I); and for the traits adjusted to variation in GY under WL05 (J), WW05 (K) and WW07 (L).

PCA conducted for all 11 elements showed that the first two PCs accounted collectively for 57%, 52.7% and 52.5% of the variations for WL05, WW05 and WW07, respectively (Figure S2). A clear clustering of three groups was shown for the first year’s data: (i) essential microelements (Cu, Fe and Zn) with GPC and S; (ii) P and Mg with Mn; and (iii) K and Ca with the non-essential Al. The WW07 PCA based on element concentrations s showed strong differences from the WWO5 data, with Al and K located outside of the group of all other elements. The first two main components of the adjusted traits PCA explained around 50% of the variation. PCA for GY- and S-adjusted traits showed a clustering pattern similar to that of the initial traits in WL05 and WW05. On the contrary, PCA for S-adjusted traits under WW07 showed clear clustering into two groups, where the first group contained Cu, Fe, Mn and Zn, and the second group included Mg, P, GPC and Al, while Ca and K showed lower associations with the other elements. PCA for P-adjusted traits showed the strongest change in clustering compared to those of the initial traits, with more clear environmental differences in the clusters. The major difference for P-adjusted traits is that K was loaded aside from the other elements. PCA of P-adjusted traits under WL05 showed the following clustering: (i) Fe, Cu and GPC; (ii) Mg, Al and Mn; and (iii) Zn and S, with Ca loading outgroup. Different associations were observed for these traits under WW05: (i) Zn, Fe and GPC; (ii) Cu and S; (iii) Mg and Mn; and (iv) Ca and Al. Three clusters can be seen for the adjusted traits in WW07: (i) Ca and Al; (ii) Cu, Fe, Mn and Zn; and (iii) Mg, GPC and S.

The effect of the adjustment of the RIL phenotypic variation in the ionome traits s was assessed using rank correlation (Kendall’s Tau) between the initial and the adjusted trait values (Table S12). High correlation implies low adjustment × genotype interaction and vice versa. Lower interaction of all adjustments were observed for the adjusted Al, Ca and Mn concentrations and higher effects were observed for GPC, Mg and Zn.

### Phenotypic association of grain ionome with other complex traits

The relationships between the ionome and a set of complex traits were characterized by estimation of correlations between the 11 elements and 17 complex traits representing yield, morphology, phenology and physiology, obtained under WL05 and WW05 (Peleg *et al.*, 2009b; Fatiukha *et al.*, 2019). Most of the ionome traits, except Mn, showed negative associations with yield, biomass related traits, and CL, with stronger negative associations obtained under WL05 (Figure 2, Table S12). SpL and leaf morphology traits (FLW and FLL) showed weak positive associations (0.17-0.28) with some of the ionome traits under WW05. Stronger associations (0.32-0.33) of SpL with Cu and Fe were observed under WL05. In contrast to WW05, associations of some of the ionome elements with FLL under WL05 were negative. Most of the physiological traits showed weak correlations with concentrations of elements under both, WW and WL, conditions. Phenological traits showed “trade of” in associations with some of the elemental traits: DP-H was positively and DH-M negatively associated with the ionome traits, with strongest correlations observed under WW05. Correlations of traits adjusted to GY showed considerable reduction of the negative associations with most of the yield related traits, whereas associations between other complex traits remained approximately the same (Figure 2).

### Genetic architecture of initial and adjusted traits

The strong interdependence between most of the elements, both with productivity and chelators, required adjustment of these traits prior to QTL analysis to avoid biases due to unaccounted phenotypic variation caused by these factors. In total, four sets of traits were used for QTL analysis: one set of 11 initial traits and three sets of traits adjusted for variation in GY, P or S concentrations. QTL analysis of all four sets of traits revealed 437 QTL effects distributed along 105 QT loci (Figure 4, Table 2, Table S13-S14). Out of the 105 QTLs, 68 exhibited pleiotropic effects on two or more traits, while37 of them affected only one trait (co-location of effects on the initial and the adjusted traits was considered as a single-trait QTL). A total of 33 QTLs affected only the adjusted traits, whereas the other 72 QTLs represented combinations of effects on the initial and adjusted traits, of which 38 QTLs displayed pleiotropic effects on the adjusted traits, despite of no effect on the corresponding initial trait.

**Fig. 4.**
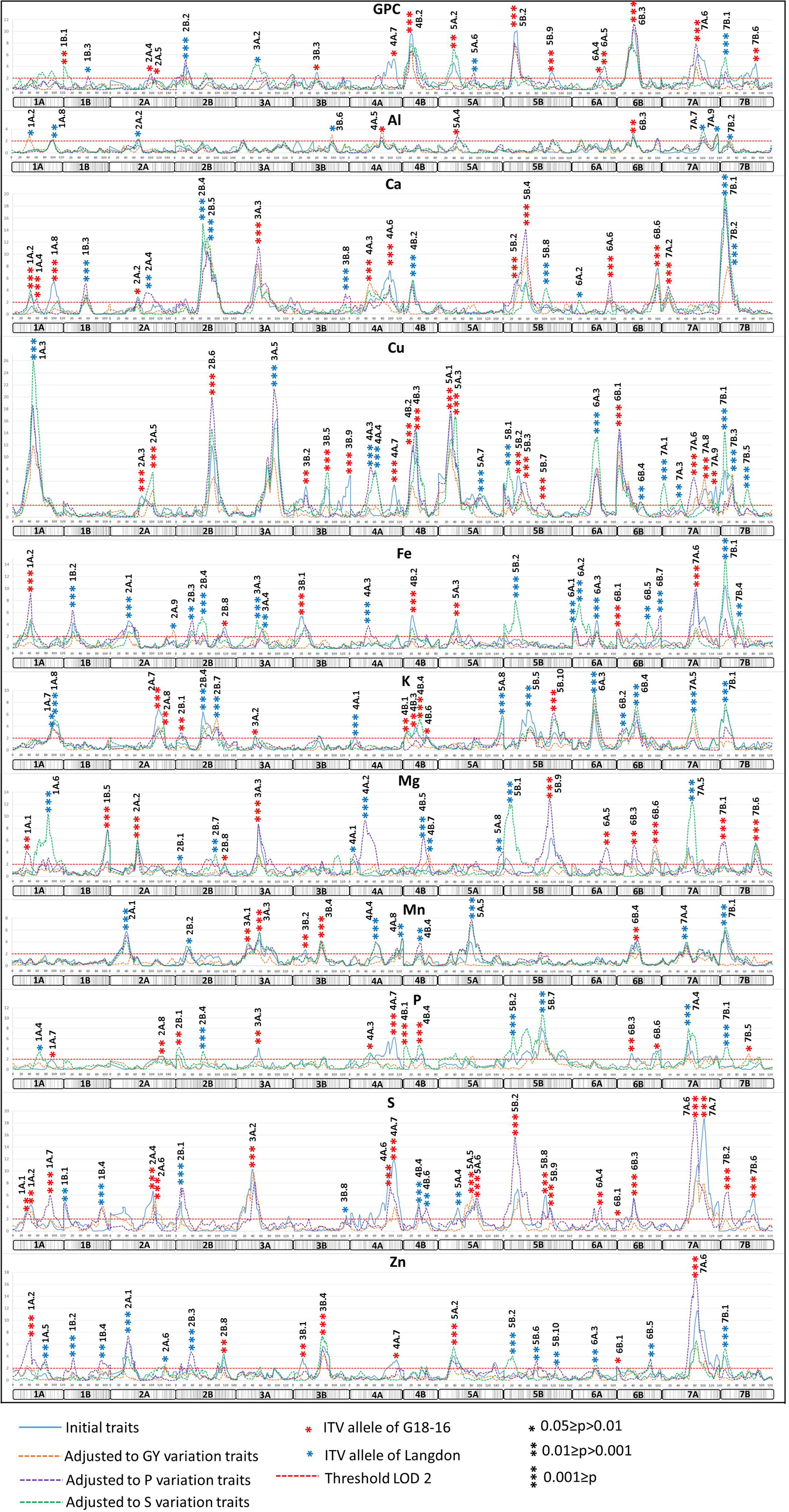
Genomic architecture of wheat grain ionome. LOD scores, significance of QTL models, increased trait value of QTL alleles and the detected QTLs are shown for 11 initial and three sets of grain ionome traits adjusted to variations in GY, P and S for the 14 chromosomes of tetraploid wheat.

The presence/absence of QTL effects on the adjusted traits were used for classification of QTLs in relation to the “adjusting factor’, i.e. GY, P or S (summarized in Table 1). In case of co-localization of the initial and the adjusted QTL effects, the effect of the QTL was referred to as “independent” from the corresponding factor. In case of the absence of adjusted QTL effects, when an effect on the initial trait was detected, QTL was classified as “dependent” on the corresponding adjusting factor. The detected QTLs were referred to as “hidden” in cases of detection of significant effects on the adjusted traits only.

**Table 1.**
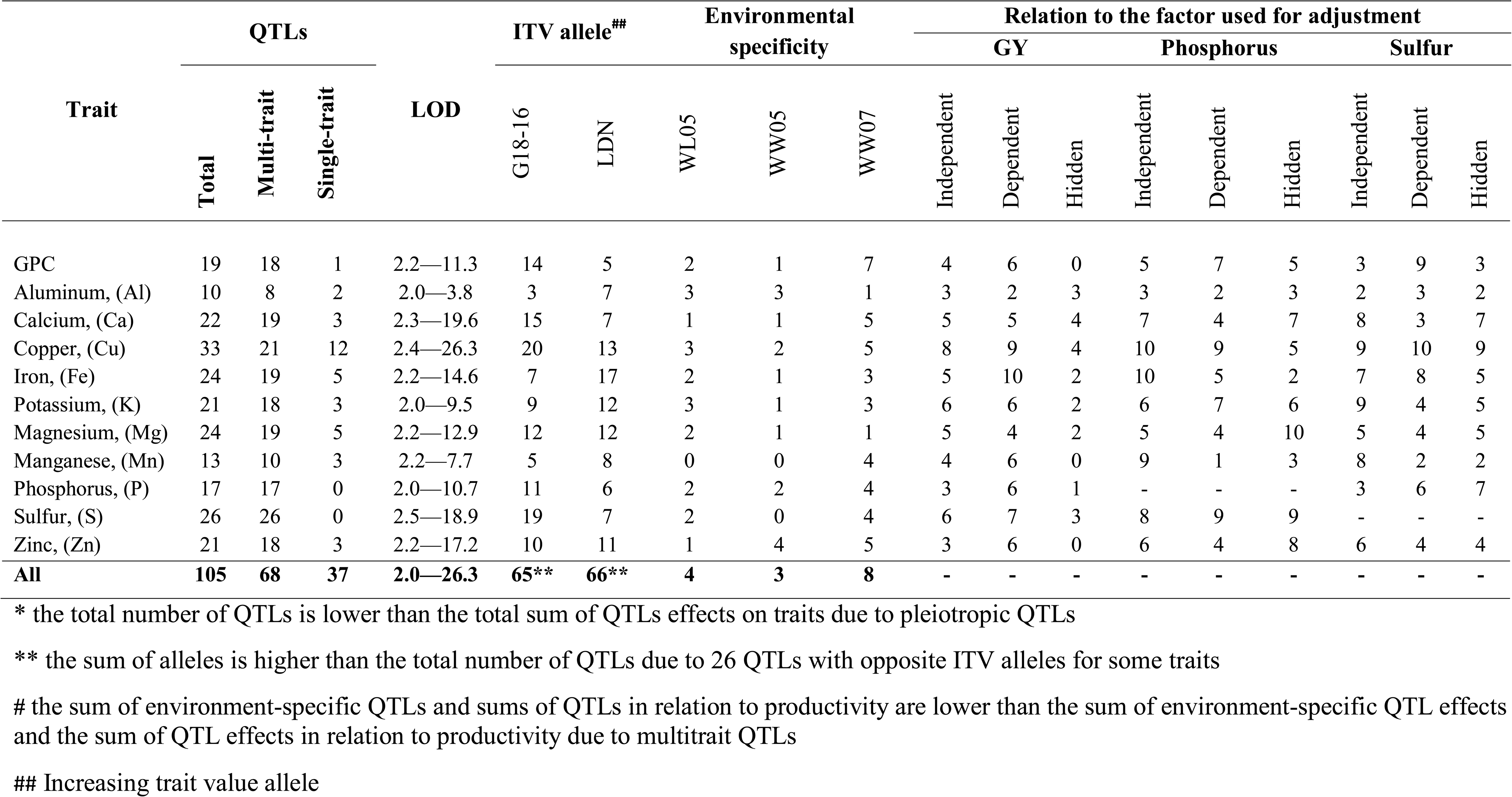
Summary of QTLs associated with grain nutrient content traits, under water limited (WL) and well-watered (WW05 and WW07) conditions.

**Table 2.** A list of selected CGresiding within the detected QTL regions for 11 grain ionome traits. IDs and Annotation of CGs is according to WEW reference genome (Avni et al, 2017).

| Traits | Chr. | QTLs | IDs of candidate genes | Annotated function |
| --- | --- | --- | --- | --- |
| GPC | 4B | 4B.2 | TRIDC4BG004570, TRIDC4BG004920 | Protein NRT1/ PTR FAMILY 8.3, Protein NRT1/ PTR FAMILY 2.11 |
|  | 5B | 5B.2 | TRIDC5BG006450, TRIDC5BG006510 | Protein NRT1/ PTR FAMILY 2.11 |
|  | 6B | 6B.3 | TRIDC6BG019590 | NAC domain protein (Gpc-B1) |
|  | 7A | 7A.6 | TRIDC7AG059770 | High affinity nitrate transporter 2.7 |
| Aluminum,<br>(Al) | 1A | 1A.2 | TRIDC1AG019270 | Calcium-transporting ATPase |
|  |  | 1A.8 | TRIDC1AG057390 | PLANT CADMIUM RESISTANCE 2 |
| Calcium,<br>(Ca) | 3A | 3A.3 | TRIDC3AG040050 | Two pore calcium channel protein 1 |
|  | 5B | 5B.4 | TRIDC5BG045550 | calcium-transporting ATPase, putative |
|  | 7B | 7B.1 | TRIDC7BG001820 | cation calcium exchanger 4 |
| Copper,<br>(Cu) | 1A | 1A.3 | TRIDC1AG037690, TRIDC1AG038820 | Cation-transporting ATPase 5, Heavy metal transport/detoxification superfamily protein |
|  | 2B | 2B.6 | TRIDC2BG062440 | Copper-transporting ATPase 1 |
|  | 5A | 5A.1 | TRIDC5AG004320 | Copper-transporting ATPase PAA1, chloroplastic |
| Iron,<br>(Fe) | 1A | 1A.2 | TRIDC1AG028130, TRIDC1AG026830 | Heavy metal transport/detoxification superfamily protein |
|  | 6A | 6A.2 | TRIDC6AG005140 | Heavy metal transport/detoxification superfamily protein |
|  | 7A | 7A.6 | TRIDC7AG058580 | vacuolar iron transporter 1 |
| Potassium,<br>(K) | 2B | 2B.4 | TRIDC2BG048260 | Potassium channel SKOR |
|  | 6A | 6A.3 | TRIDC6AG029740, TRIDC6AG044480 | Potassium transporter family protein |
|  | 6B | 6B.4 | TRIDC6BG028640 | Potassium channel KAT1 |
|  | 7A | 7A.5 | TRIDC7AG031680, TRIDC7AG050040 | Potassium transporter 26, Glutathione-regulated potassium-efflux system protein |
| Magnesium,<br>(Mg) | 2A | 2A.2 | TRIDC2AG032490 | Magnesium transporter MRS2-B |
|  | 4A | 4A.3 | TRIDC4AG044250 | Magnesium transporter MRS2-B |
|  | 7A | 7A.5 | TRIDC7AG031890, TRIDC7AG047050 | Heavy metal transport/detoxification superfamily protein |
| Manganese,<br>(Mn) | 2A | 2A.1 | TRIDC2AG011890, TRIDC2AG011900 | NAC domain protein |
|  | 3A | 2A.1 | TRIDC3AG027860, TRIDC3AG030720 | Divalent metal cation transporter MntH, Heavy metal transport/detoxification superfamily protein |
|  | 5A | 5A.5 | TRIDC5AG049910 | NAC domain protein |
| Phosphorus,<br>(P) | 4A | 4A.7 | TRIDC4AG062110, TRIDC4AG062150 | phosphate transporter 1;5, phosphate transporter 1;1 |
|  | 4B | 4B.1 | TRIDC4BG001410, TRIDC4BG002220 | phosphate transporter 4;1, phosphate transporter 1;1 |
|  | 7A | 7A.4 | TRIDC7AG022170, TRIDC7AG022180 | Phosphate-responsive 1 family protein |
| Sulfur,<br>(S) | 3A | 3A.2 | TRIDC3AG010550 | cysteine synthase D1 |
|  | 4A | 4A.7 | TRIDC4AG058580 | sulfate transporter 3;4 |
|  | 7A | 7A.6 | TRIDC7AG052360, TRIDC7AG054490 | Glutathione S-transferase family protein, glutamine dumper 4 |
| Zinc,<br>(Zn) | 1A | 1A.2 | TRIDC1AG014690, TRIDC1AG019100 | Zinc transporter 6, Zinc transporter ZupT |
|  | 3B | 3B.4 | TRIDC3BG057740 | Metal tolerance protein 5 |
|  | 7A | 7A.6 | TRIDC7AG058500 | Zinc transporter 10 |

Mapping of the adjusted traits considerably improved the QTL detection power enabling identification of new QTL effects compared to those detected on initial traits (Figure 4-6, Table S13). For example, we detected 24 QTLs with effect on Mg concentration. Out of them, 15 QTLs were found for the adjusted traits only (Figure 6), 5 QTLs showed an increase in LOD score for adjusted traits compared to that of the initial traits (Figure 5), and the remaining 4 QTLs displayed a decrease of the LODs for the adjusted traits. For Cu concentration, 14 QTLs out of 33 were detected for adjusted traits only (Figure 6), LOD scores were increased for 11 QTLs (Figure 5) and decreased for 8 adjusted traits.

**Fig. 5.**
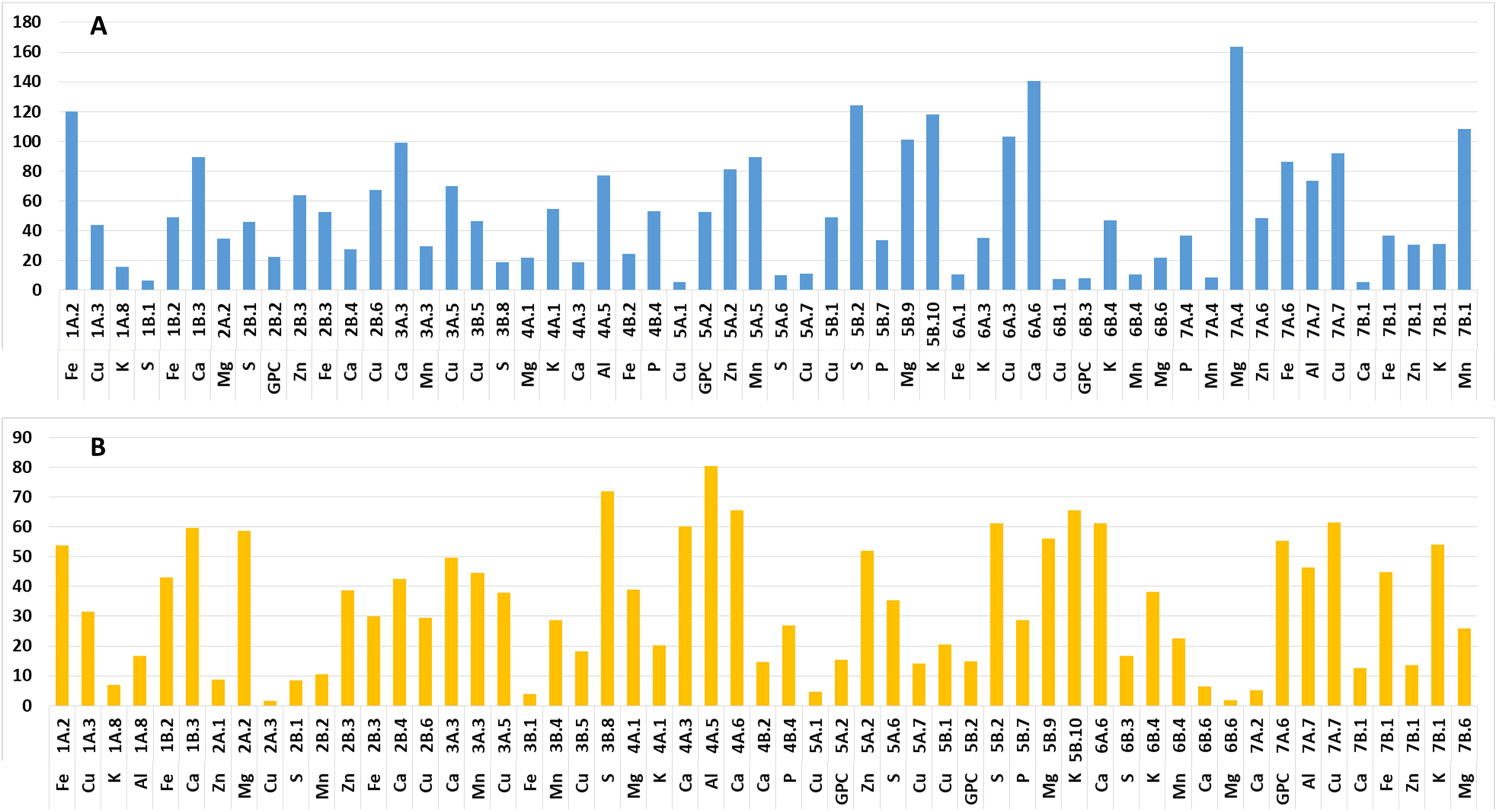
Improvements of LOD scores and confidential intervals (represented by1.5 LOD support intervals) of QTL effects for the adjusted traits compared to the intial traits. (A) Percent of increasing of LOD scores: ΔLOD=100· (LOD_adjusted_-LOD_initial_)/LOD_initial_. (B) Percent of decreasing of confidential intervals (CI) ΔCI=100·(CI_adjusted_-CI_initial_)/CI_initial_.

**Fig. 6.**
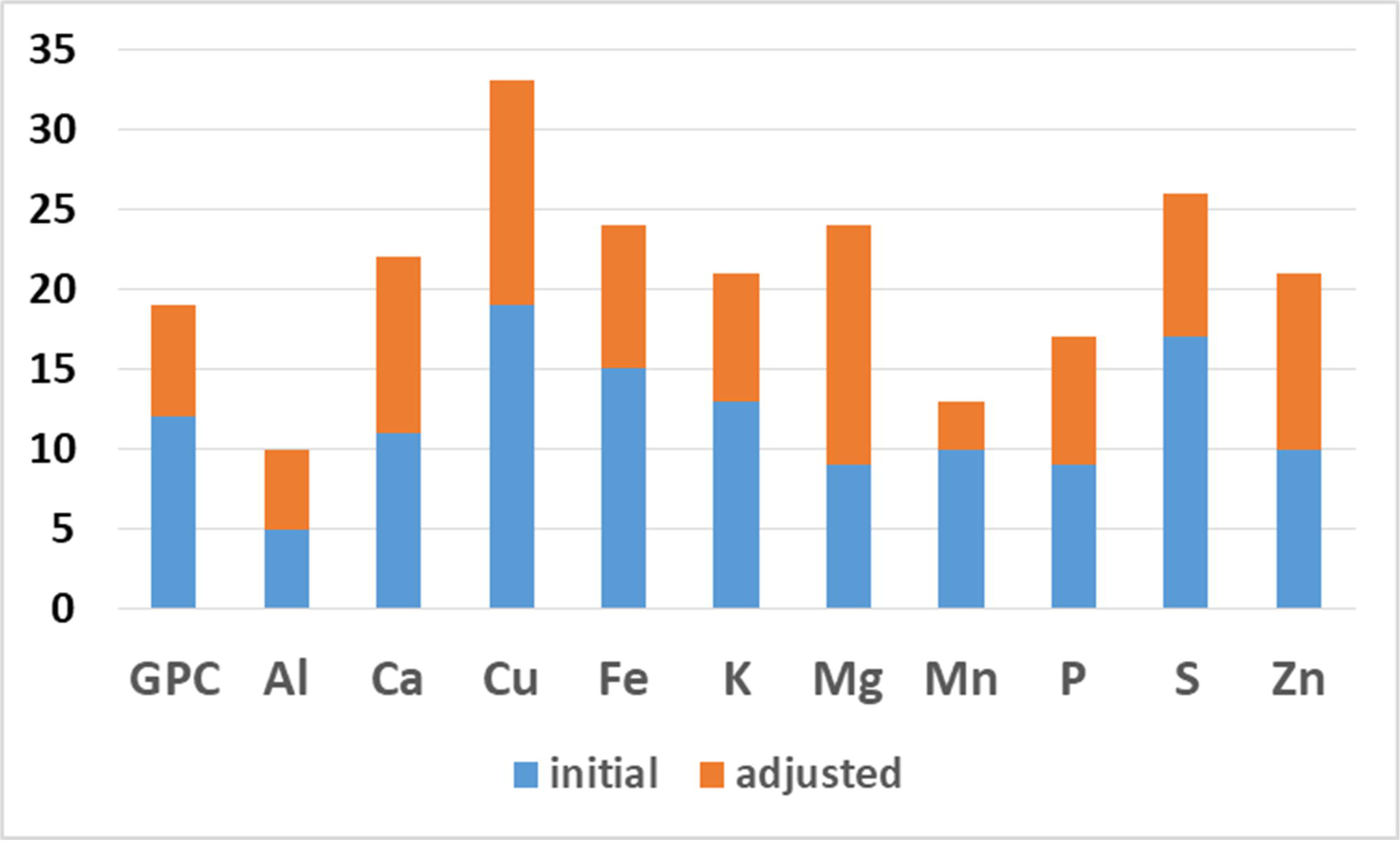
Total number of QTL effects identified for 11 elemental ionome traits. QTL effects for the adjusted traits were counted in this illustration only in cases of no significant effect on the corresponding initial trait.

Since one parent of the studied RIL population was WEW, it was interesting to compare positive effects on concentrations of elements conferred by cultivar vs. wild alleles (Figure 7). A clear advantage of WEW as a source of positive alleles was shown for GPC and S (see Fig. 7) indicating that the employed WEW genotype can be considered as a promising source for increasing protein and S content. The advantage of WEW for other traits (Cu, Mg and Zn) is less preannounced and may vary upon comparisons to other cultivated genotypes.

**Fig. 7.**
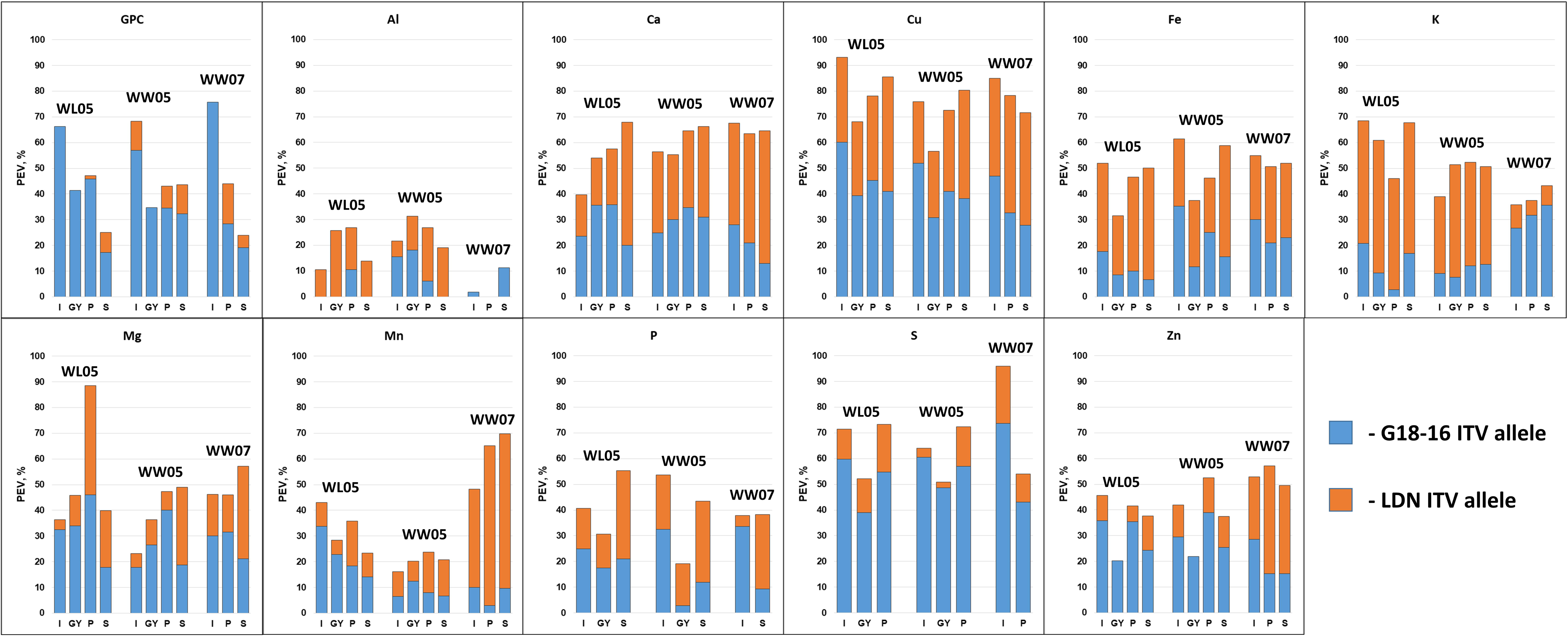
Comparisons of the parental ITV allele contribution in the genetic regulation of 11 grain ionome traits detected in RIL population grown under three environments. These comparisons are based on percent of explained variation of the detected QTL effects.

### Shared genetic basis of wheat grain ionome

We have ranked the ionome elements according to the total number of co-localized QTL effects with other traits: S (58) > Fe (51) > Ca (48) = Cu (48) > P (47) > Zn (43) > GPC (42) > Mg (38) > K (37) > Mn (24) > Al (18) (Table S15). The maximum number of common QTLs, providing an indication for a shared regulation, was found for Fe/Zn (12), and GPC/S (11). On average, each element has about four common QTLs with other elements, while only Mn and Al exhibited no shared QTL effects (Table S16). The non-essential element Al showed the weakest QTL co-localization with other elements (on average 1.6), and Mn was found as second in weak co-regulation with other element (2.2 common QTLs on average). The adjustment of element concentrations for variation in yield, P, and S, allowed us to improve the QTL detectability and the mapping accuracy, leading to more objective conclusions about co-localization of QTL effects for different traits. These adjustments resulted in the detection of 24 additional common QTL effects, on average, for each element, with the lowest number for Mn (10) and the highest number for S (36) (Table S16).

We have compared the obtained results on common QTLs for the different ionome elements also with the phenotypic associations of these elements, in the RIL population, based on correlation analysis (described in section Phenotypic association within wheat grain ionome) (Table S17). For most of the combinations, we found rather strong correspondence between the level of co-localization and phenotypic associations. However, some of the combinations showed unbalanced ranks of phenotypic associations with the obtained genetic architecture (Table S17), which can be explained by the presence of QTLs with opposite increasing trait value (ITV) alleles or differences in the effects (e.g. percent of explained variation, PEV) for the associated traits (Table S15). Yet, some of the elements showed only a minor level of co-localization of QTL effects, despite the rather strong phenotypic associations, as demonstrated by Mg-P and GPC-P.

### Candidate genes

In our previous study (Fatiukha *et al.*, 2019), we have anchored most of the SNPs used in the current QTL mapping to the annotated reference genome of WEW (Avni *et al.*, 2017). Thus, we were able to estimate the physical intervals of the detected QTLs using the relative proportion between the physical and genetic positions of those SNPs and the 1.5 LOD support interval of the detected QTLs. The physical interval lengths of the QTLs ranged from 4.4 to 455.6 Mbp with the number of genes within these intervals varying from 25 to 1740 (Tables S14, S18). The distribution of the physical lengths of QTLs showed a clear partition into two groups: (i) up to 150 Mbp (with sub-telomeric and mid-arm locations), and (ii) longer than 250 Mbp (located mostly in pericentromeric regions). According to the gene models described in Avni *et al.*, (2017), the support intervals of 51 out of the 105 detected QTLs included less than 200 high confidence (HC) genes per interval, therefore, allowing to conduct a search for the underlying CGs.

We have focused on genes involved in transport and metabolism of ions and elements as potential CGs and selected a total of 691 CGs within all of the QTL intervals (Table S19). We have identified some strong associations of CG with specific function that directly matched the established QTL effects (Table 2). For example, three QTLs (3A.3, 5B.4 and 7B.1) with strong effects on Ca concentration (LOD scores 11.2, 14.6 and 20.6, respectively) included CG related to Ca metabolism: *TRIDC3AG040050* (two pore calcium channel protein 1), *TRIDC5BG045550* (calcium-transporting ATPase), and *TRIDC7BG001820* (cation calcium exchanger 4). Strong associations for Cu (2B.6, 5A.1, and 7A.6 with LODs 20.1, 17.7, and 6.6, respectively) and Fe (1A.2, 2B.4, and 7A.6 with LODs 9.2, 5.2, and 10.0, respectively) included the following genes: *TRIDC2BG062440* (copper-transporting ATPase 1), *TRIDC5AG004320* (copper-transporting ATPase PAA), *TRIDC7AG058420* (copper-transporting ATPase 1), *TRIDC1AG015400* (4Fe-4S ferredoxin, iron-sulfur binding protein), *TRIDC2BG050320* and *TRIDC7AG058580* (vacuolar iron transporter 1). The strongest QTL affecting K (6A.3 with LOD 9.5) included three genes related to the potassium transporter family (*TRIDC6AG029740, TRIDC6AG044480*, and *TRIDC6AG044820*).

In addition, we found a few families of genes that were more frequently present within QTL intervals than representatives of other families. For example, we identified 77 genes related to heavy metal transport/detoxification superfamily proteins; 14 genes associated with metal tolerance; 43 genes related to the NAC domain containing proteins; and 46 genes related to the NRT1/PTR family. Although some of the QTL effects had strong functional associations with the identified CGs (Table 2), families of genes that are associated with a specific element can be involved also in metabolism of other elements due to the similarity in the chemical properties. For example, we have identified 33 and 23 genes involved in Zn and Cu transport, respectively; similarly, 32 and 26 genes were associated with Ca and Mg transport, respectively. Identification of CGs associated with P and S metabolism was limited due to the complexity of their genetic regulations; thus, such candidate genes can be identified in almost each QTL interval. In total, we have identified 39 CGs associated with phosphate transport, 47 glutathione CGs, and 16 CGs associated with sulfate transport. The well-known gene *Gpc-B1*, which is involved in remobilization of proteins and micronutrients in wheat (Uauy et al. 2006), was found in the 6B.3 GPC QTL interval. In addition, *Vrn-B3* was found to reside in the the 7B.1 QTL, which showed effects on eight ionome traits, making it a strong CG conferring these traits.

## Discussion

The recent plant ionomics studies show a substantial progress in the understanding of the genetic regulations of elemental metabolism (Huang & Salt, 2016). Although rich ionome data is now available for different plant species (Baxter *et al.*, 2007), the extent of ionome studies in wheat is still limited. A major advance in plant iomonics was achieved by the analysis of the elemental composition as a network that enables to dissect the relationships and interdependence between the elements (Baxter, 2015). Some of the associations between elements can be explained by the common biochemical nature and biological function they share in plants (Pauli *et al.*, 2018). Yet, other elements, also showing strong phenotypic associations, for example, between P and Mg in cereal grains (Shakoor *et al.*, 2016), cannot be explained using this hypothesis due to their different chemical nature and metabolic roles in plants. Overlooking these differences can lead to wrong interpretations, where, for example, association between two elements can be induced by a third one. Furthermore, when applying genetic analyses, such as QTL and GWAS, this can lead to biases in mapping due to incorrect (biased) phenotypes. The working hypothesis of our study was based on the assumption that two major structural elements of chelators, P and S, shape the elemental composition of wheat grain ionome. Although variation in chelate accumulation was proposed earlier as an important factor that can alter element level in plants (Baxter, 2009), we did not encounter studies that implement this idea in genetic analysis. In this study, our main goal was to dissect the genetic architecture of wheat grain ionome. Thus, we conducted QTL analysis of both, the initial elemental grain concentrations, as well as of the traits adjusted for P and S, in order to reduce the biases caused by their phenotypic associations with other elements. Furthermore, we adjusted the element concentrations for the variation in productivity in order to avoid the “dilution effect” (Davis, 2009), as another source of biases.

The approach of adjusted traits based on residuals from regression was previously demonstrated for accounting variation in weight of grains in ionome studies (Shakoor *et al.*, 2016), as well as for accounting for the negative correlations commonly observed between yield and GPC (Oury & Godin, 2007).. In the current study, the adjustments for variation in P and S showed considerable effects on the phenotypic associations, while accounting for the “dilution effect” did not show obvious effects on the associations within ionome. Since phytates represent 60-80% of the total P content in seeds (Lott *et al.*, 2000) and S compounds are intensely involved in metal chelation (Na & Salt, 2011), we used P and S concentrations as estimators for the corresponding chelators. Although, S concentration might be considered as a weak chelate estimator in comparison to P, since a high proportion of S is stored in grains as a part of storage proteins (Galili, 1997), nevertheless, a considerable improvement in QTL detectability was observed after S adjustment. Obviously, the use of direct measurement of chelators would provide a better accounting for their effects, however, practically, it cannot replace ionomics as a high-throughput phenotypic approach (Baxter, 2010). Although our approach seems to be efficient in accounting for P and S, it cannot be used for adjusting the variation in N containing chelators (Leitenmaier & Küpper, 2013), since most of the N is stored in grain storage proteins.

The phenotypic associations, observed in our study, corroborate well with what we know about the biochemical composition of cereal grains and are in agreement with other studies of phenotypic associations within grain ionome of major cereals, such as sorghum (Shakoor *et al.*, 2016), maize (Baxter *et al.*, 2013; Gu *et al.*, 2015), rice (Stangoulis *et al.*, 2007), and wheat (Morgounov *et al.*, 2007; Gomez-Becerra *et al.*, 2010b; Pandey *et al.*, 2016). Most of these studies reported positive associations of the essential metal microelements- Cu-Fe-Zn with GPC and S, and the essential metal macroelements Ca-K-Mg with P in cereal grains. Phytates serve as a major storage for Mg, Ca and K in seeds (Lott *et al.*, 2000), although other metals also can be stored in combination with phytic acid (Xue *et al.*, 2015), especially in wheat (Ficco *et al.*, 2009). Moreover, we can assume that some negative associations of K after adjustment for P concentration can result from the competition between divalent Mg and Ca with monovalent K (Lott *et al.*, 1985). It is also important to mention that in cereal crops, most of the plant K (over 70 %) is accumulated in the straw and less K is allocated to grain (Zörb *et al.*, 2014), thus adding another factor that may contribute to the mentioned negative association. The strong phenotypic correlation between GPC and P is probably associated with the balance between the macroelements needed for the globoid crystal formations (Lott *et al.*, 1985), where storage proteins may be associated with chelate complexes (Raboy *et al.*, 1991). In the current study, we have obtained strong associations between S and GPC, which can be explained by the strong coordination of N and S metabolisms in wheat that are affecting amino acid metabolism (Howarth *et al.*, 2008) and grain proteome (Dai *et al.*, 2015; Bonnot *et al.*, 2017). Moreover, similar linkage was observed in barley when N deprivation affected also cysteine biosynthesis (Carfagna *et al.*, 2011). The strong pleiotropic/linkage QTL effects on S and some metals observed in the current study are supported by previous publications showing that genes involved in the metabolism of S compounds affected also metal accumulation. For example, expression of wheat phytochelatin synthase gene (*TaPCS1*) in rice resulted in increased Cd sensitivity (Wang *et al.*, 2012); metallothionein related genes influenced Cu accumulation in Arabidopsis (Benatti *et al.*, 2014) and zinc accumulation in barley (Hegelund *et al.*, 2012). In addition, S nutrition level can affect Fe accumulation in durum wheat seeds (Astolfi *et al.*, 2018). The strong genetic associations between Fe and Zn, previously shown also in other studies (Cakmak *et al.*, 2010), can be explained by the shared transporters (Kobayashi & Nishizawa, 2012).

The elemental spatial distribution maps within cereal grains, obtained from X-ray fluorescence microscopy provide another layer of support for the consideration of P and S concentrations as estimators of chelators. Most of the published maps showed co-localization of P and S with Cu, Fe, Zn and Mn in barley and wheat grains (Lombi *et al.*, 2011; Ajiboye *et al.*, 2015; De Brier *et al.*, 2016). Although, element- and tissue-specific localizations were found for different metals, the aleurone layer showed the highest co-localization of P and S with metals. In contrast, some barley genotypes showed higher accumulation of Zn in the endosperm where it can create a complex with S ligands (Detterbeck *et al.*, 2016). The N and S nutrition levels have shown strong effects on the deposition and spatial distribution of Zn in the endosperm of wheat grain (Persson *et al.*, 2016); similar results were obtained for biofortified wheat grains under different nutrient concentrations (Ramos *et al.*, 2016).

Ionome relationships with other complex traits, such as morphology and phenology, can alter the concentrations of elements. However, according to our data most of the other complex traits showed weak environment- and trait-specific associations with element concentrations. Only yield related traits showed a common pattern of negative association with ionome traits. Therefore, we selected GY as an integrative dilution factor that can account for the effects of most of the productivity components. Although, adjustment of variation in GY mostly did not affect phenotypic association as well as genetic architecture, the obtained results of QTL classification in relation to productivity can serve as a source of selection of more promising QTL alleles for marker assistant breeding.

Although the improvement of the nutritional quality of staple crops is an important agronomic challenge (White & Broadley, 2009), most of the studies dealing with the genetic analyses of elemental composition in wheat are focused on Fe and Zn (Borrill *et al.*, 2014) as the major malnutrition factors (Younis *et al.*, 2015). Some studies showed strong effect of the metabolism of the structural elements (N, P and S) on plant ionome (Na & Salt, 2011; Ding *et al.*, 2017), accounting for these effects in biofortification programs is limited and thus our approach can improve their efficiency. WEW was previously shown as a source for improvement of nutrition quality of modern wheat (Gomez-Becerra *et al.*, 2010a; Chatzav *et al.*, 2010; Klymiuk et al. 2019). Our results also show the potential of using WEW beneficial alleles for increasing of GPC and S grain concentrations. This is highly valuable for wheat breeding since S metabolism plays a crucial role in S-rich and S-poor storage proteins that strongly affect wheat baking quality (Shewry *et al.*, 2001).

Identification of CGs involved in regulation of elemental accumulations is an important part of grain ionomics. While QTL analysis cannot provide resolution at the level of a single gene, recent advantages in wheat genomics (Avni *et al.*, 2017; Appels *et al.*, 2018) allowed us to directly translate genetic intervals to sequence intervals on wheat pseudomolecules and identify CGs within chromosome segments that represent these QTL intervals. These results can serve as a solid base for future works combining transcriptome studies, allele mining and even cloning of the strong CGs. We believe that despite the relatively large number of genes that reside within ionome QTL intervals, the functional annotations of the CGs can be used for identification of the most promising ones, as demonstrated in Table 2. Obviously, the limitations caused by insufficient mapping resolution of QTL analysis may be considerably relaxed by applying our “adjustment” approach in GWAS. Following the idea of shared genetic regulation of metals, we have identified numerous heavy metal transport/detoxification superfamily genes within the 41 QTL intervals. This family is involved in the regulation of the accumulation of many metals, including the essential ones (de Abreu-Neto *et al.*, 2013). An obvious CG for the major GPC QTL identified in the current study on chromosome arm 6BS is *Gpc-B1* that belongs to the NAC family of transcription factors (TFs), and was shown to be involved in the regulation of nutrient remobilization (Uauy *et al.*, 2006). Genes related to glutathione S-transferase family were shown to be involved in Cu tolerance in rice (Li *et al.*, 2018) and were identified in the current study within 25 QTL intervals, therefore, supporting our hypothesis that S metabolism is associated with the genetic regulation of metal accumulation.

## Conclusions

The functional basis for the physiological, biochemical and genetic regulations of accumulation, transport, and up-take of elements in plants is an important target for fundamental and applied research studies. In this work, we have demonstrated the influence of the variation in P and S concentrations, as estimators of the corresponding chelators, on the phenotypic associations, as well as on the genetic architecture of grain ionome. We have applied a new approach for adjustment of these effects in genetic analysis that resulted in considerable increase of the QTL detection power leading to the identification of new ionome QTLs. The obtained results showed improved correspondence between the genetic basis and the phenotypic associations, confirming the importance of understanding of the interdependence of the ionome. Our approach can be utilized in a wide range of physiological and genetic studies that are focused on the regulation of ionome or certain elements. The identification of CGs within QTL intervals provides an important basis for future studies on the genetic and molecular mechanisms controlling wheat grain ionome. In addition, the identified WEW QTL alleles leading to increase in GPC and essential elements, such as Zn and Fe, emphasize the potential of wild relatives for improvement of nutrition quality of crops.

## Acknowledgements

The research leading to these results has received funding from the European Community’s Seventh Framework Programme (FP7/ 2007-2013) under the grant agreement n°FP7-613556, Whealbi project; the Israeli Ministry of Agriculture and Rural Development, Chief Scientist Foundation (Grants 837-0079-10 and 837-0162-14); the US-Israel Binational Agricultural Research & Development Fund (US-4916-16); and ISF grant for equipment (grant no. 2289/16 and 2342/18), YS is the incumbent of the Haim Gvati Chair in Agriculture.

## Author Contribution

A.F., A.B.K., T.K, Y.S. and T.F. designed the research; A.F. and V.K. performed the data analysis; A.F performed QTL analysis; and Z.P. performed field phenotyping; I.C. performed elemental analysis, A.F., V.K., A.B.K., and T.F. wrote the manuscript.

## Supporting Information

Additional supporting information can be found in the online version of this article.

Fig. S1 Phenotypic distribution of 11 ionome traits under three environments (WL05, WW05 and WW07) in G×L RIL population.

Fig. S2 Principal Component Analysis (PCA) of the initial and the adjusted (to variation in GY, P and S) ionome traits obtained for the G×L RIL population under three environments WL05, WW05 and WW07.

Table S1 Analyses of variance (Anova), means and ranges for the 11 ionome traits.

Table S2 Ranges of the adjusted ionome traits to variation in GY, P and S under the three environments WL05, WW05 and WW07.

Table S3 Coefficients of correlation (r) between ionome traits under WL05.

Table S4 Coefficients of correlation (r) between ionome traits under WW05.

Table S5 Coefficients of correlation (r) between ionome traits under WW07.

Table S6 Coefficients of correlation (r) between the adjusted ionome traits to variation in GY under Wl05 and WW05.

Table S7 Coefficients of correlation (r) between the adjusted ionome traits to variation in GY under WW07.

Table S8 Coefficients of correlation (r) between the adjusted ionome traits to variation in P and coefficients of correlation (r) between the adjusted ionome traits to variation S under WL05.

Table S9 Coefficients of correlation (r) between the adjusted ionome traits to variation in P and coefficients of correlation (r) between the adjusted ionome traits to variation S under WW05.

Table S10 Coefficients of correlation (r) between the adjusted ionome traits to variation in P and coefficients of correlation (r) between the adjusted ionome traits to variation S under WW07.

Table S11 Kendall’s tau coefficients of rank correlations between the initial and the adjusted traits under the three environments WL05, WW05 and WW07.

Table S12 Coefficients of correlation (r) between 17 initial traits and ionome traits in two treatments (WL05 and WW05) and GY in WW07.

Table S13 Parameters of QTL effects for the 11 initial ionome traits and the adjusted to variation in GY, P and S traits in G×L RIL population under the three environments WL05, WW05 and WW07.

Table S14 Summary of QTLs detected for the initial ionome traits and the adjusted to variation in GY, P and S in G×L RIL population under the three environments (WL05, WW05 and WW07).

Table S15 Genetic associations between the ionome traits based on the total number of common QTLs.

Table S16 Number of QTLs detected for the initial and the adjusted traits that share co-localization with other traits.

Table S17 Comparisons of genetic and phenotypic associations between the ionome traits.

Table S18 A list of HC genes within QTL intervals.

Table S19 Summary of selected CGs.

