## Supplementary Data 1 for "Variation in phosphorus and sulfur content shapes the genetic architecture and phenotypic associations within wheat grain ionome"

**Fig. S1** Phenotypic distribution of 11 ionome traits under three environments (WL05, WW05 and WW07) in G×L RIL population.

**Fig. S2** Principal Component Analysis (PCA) of the initial and the adjusted (to variation in GY, P and S) ionome traits obtained for the G×L RIL population under three environments WL05, WW05 and WW07.

**Table S1** Analyses of variance (Anova), means and ranges for the 11 ionome traits

**Table S2** Ranges of the adjusted ionome traits to variation in GY, P and S under the three environments WL05, WW05 and WW07.

**Table S3** Coefficients of correlation (r) between ionome traits under WL05.

**Table S4** Coefficients of correlation (r) between ionome traits under WW05.

**Table S5** Coefficients of correlation (r) between ionome traits under WW07.

**Table S6** Coefficients of correlation (r) between the adjusted ionome traits to variation in GY under Wl05 and WW05.

**Table S7** Coefficients of correlation (r) between the adjusted ionome traits to variation in GY under WW07.

**Table S8** Coefficients of correlation (r) between the adjusted ionome traits to variation in P and coefficients of correlation (r) between the adjusted ionome traits to variation S under WL05.

**Table S9** Coefficients of correlation (r) between the adjusted ionome traits to variation in P and coefficients of correlation (r) between the adjusted ionome traits to variation S under WW05.

**Table S10** Coefficients of correlation (r) between the adjusted ionome traits to variation in P and coefficients of correlation (r) between the adjusted ionome traits to variation S under WW07.

**Table S11** Kendall's tau coefficients of rank correlations between the initial and the adjusted traits under the three environments WL05, WW05 and WW07.

**Table S12** Coefficients of correlation (r) between 17 initial traits and ionome traits in two treatments (WL05 and WW05) and GY in WW07.

**Table S13** Parameters of QTL effects for the 11 initial ionome traits and the adjusted to variation in GY, P and S traits in G×L RIL population under the three environments WL05, WW05 and WW07.

**Table S14** Summary of QTLs detected for the initial ionome traits and the adjusted to variation in GY, P and S in G×L RIL population under the three environments (WL05, WW05 and WW07).

**Table S15** Genetic associations between the ionome traits based on the total number of common QTLs.

**Table S16** Number of QTLs detected for the initial and the adjusted traits that share co-localization with other traits.

**Table S17** Comparisons of genetic and phenotypic associations between the ionome traits.

**Table S18** A list of HC genes within QTL intervals.

**Table S19** Summary of selected CGs.

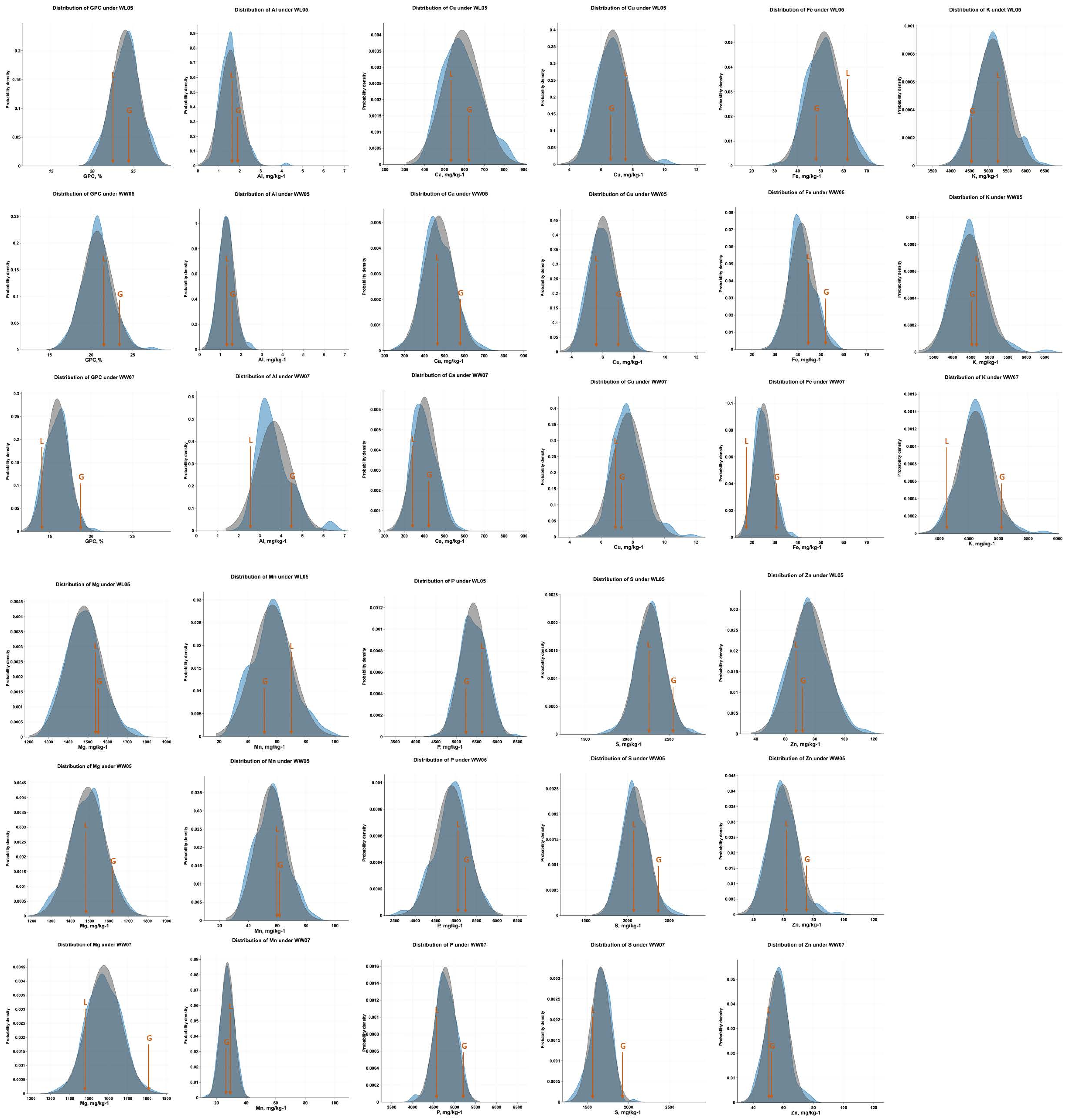


**Fig. S1.** Phenotypic distribution of 11 ionome traits under three environments (WL05, WW05 and WW07) in G×L RIL population. The arrows represent the average values of the parental lines, Langdon (L) and G18-16 (G). The expected normal distribution for each trait is presented in gray, while the observed distribution is presented in blue.

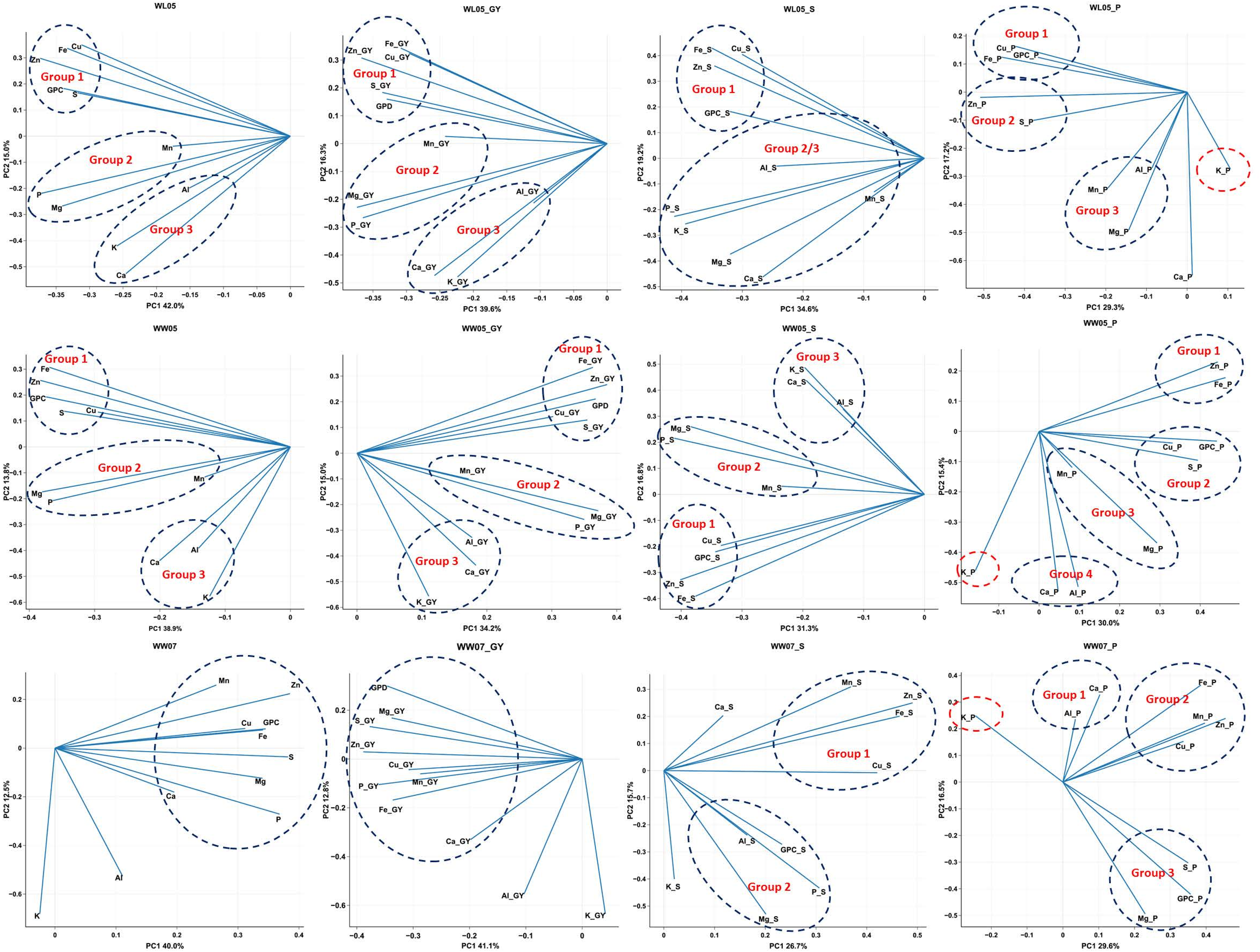


**Fig. S2**. Principal Component Analysis (PCA) of the initial and the adjusted (to variation in GY, P and S) ionome traits obtained for the G×L RIL population under three environments WL05, WW05 and WW07.

**Table S1** Analyses of variance (Anova), means and ranges for the 11 ionome traits: grain protein content (GPC), aluminum (Al), calcium (Ca), copper (Cu), iron (Fe), potassium (K), magnesium (Mg), manganese (Mn), phosphorus (P), sulfur (S), zinc (Zn) in G×L RIL population under three environments WL05, WW05 and WW07.

| | **GPC** | **Al** | **Ca** | **Cu** | **Fe** | **K** | **Mg** | **Mn** | **P** | **S** | **Zn** |
|---|---|---|---|---|---|---|---|---|---|---|---|
| **Irrigation MS (DF=1)** | 9.05<br>** | 372.10<br>*** | 12.17<br>*** | 275.20<br>*** | 28.23<br>*** | 64.53<br>*** | 20.47<br>*** | 148.98<br>*** | 37.23<br>*** | 43.72<br>*** | 39.91<br>*** |
| **Environment MS (DF=2)** | 1682.41<br>*** | 1103.14<br>*** | 483.51<br>*** | 230.97<br>*** | 1515.22<br>*** | 119.24<br>*** | 95.16<br>*** | 538.07<br>*** | 241.69<br>*** | 1184.92<br>*** | 297.72<br>*** |
| **Year MS (DF=1)** | 1877.91<br>*** | 2134.41<br>*** | 445.23<br>*** | 360.99<br>*** | 2059.86<br>*** | 39.31<br>*** | 183.76<br>*** | 1074.97<br>*** | 167.93<br>*** | 1846.40<br>*** | 205.46<br>*** |
| **WL05** | | | | | | | | | | | |
| **Mean** | 24.07 | 1.56 | 587.65 | 6.73 | 51.50 | 5114.46 | 1478.80 | 56.57 | 5414.69 | 2275.03 | 76.02 |
| **Range** | 20.06–27.90 | 0.37–4.21 | 402.87–819.51 | 4.44–10.10 | 30.80–70.27 | 4084.17–6445.85 | 1290.66–1741.60 | 30.86–96.50 | 4516.37–6414.65 | 1750.51–2774.64 | 48.91–114.42 |
| **LDN** | 23.73 | 1.72 | 532.66 | 7.38 | 61.43 | 5377.56 | 1545.20 | 67.91 | 5566.28 | 2316.48 | 69.63 |
| **G18-16** | 24.87 | 1.97 | 623.90 | 6.55 | 48.04 | 4517.98 | 1559.91 | 52.26 | 5203.96 | 2567.11 | 71.23 |
| **Genotype MS (DF=149)** | 1.96<br>*** | 1.32<br>* | 2.80<br>*** | 5.75<br>*** | 3.09<br>*** | 2.43<br>*** | 3.06<br>*** | 1.12<br>n.s. | 2.21<br>*** | 2.46<br>*** | 2.26<br>*** |
| **WW05** | | | | | | | | | | | |
| **Mean** | 20.71 | 1.29 | 471.00 | 6.01 | 41.81 | 4465.18 | 1491.05 | 55.82 | 4884.26 | 2089.94 | 59.58 |
| **Range** | 16.31–27.25 | 0.44–2.50 | 269.07–697.61 | 4.14–8.37 | 29.32–80.75 | 3525.66–6542.26 | 1264.33–1714.00 | 34.98–85.01 | 3613.95–5778.69 | 1725.33–2598.98 | 38.88–95.70 |
| **LDN** | 20.63 | 1.33 | 466.03 | 5.84 | 45.64 | 4647.31 | 1471.53 | 59.01 | 5074.62 | 2072.21 | 62.67 |
| **G18-16** | 24.13 | 1.56 | 592.42 | 7.06 | 52.63 | 4540.50 | 1628.76 | 61.70 | 5183.62 | 2399.16 | 76.92 |
| **Genotype MS (DF=149)** | 3.18<br>*** | 1.27<br>* | 4.95<br>*** | 2.18<br>*** | 2.92<br>*** | 1.60<br>*** | 2.43<br>*** | 1.34<br>* | 2.15<br>*** | 3.05<br>*** | 2.01<br>*** |
| **WW07** | | | | | | | | | | | |
| **Mean** | 15.90 | 3.65 | 400.99 | 7.70 | 25.06 | 4613.29 | 1576.13 | 27.31 | 4779.36 | 1663.28 | 55.98 |
| **Range** | 12.98–20.31 | 2.10–6.48 | 270.06–578.00 | 5.37–11.66 | 16.78–37.69 | 3926.48–5754.54 | 1333.08–1821.97 | 13.91–37.93 | 4008.15–5353.91 | 1321.57–2058.59 | 39.32–77.84 |
| **LDN** | 14.14 | 2.55 | 336.71 | 7.17 | 17.50 | 4153.10 | 1469.06 | 28.51 | 4458.02 | 1564.38 | 53.53 |
| **G18-16** | 19.48 | 4.44 | 416.00 | 7.65 | 30.11 | 5066.97 | 1805.10 | 26.91 | 5178.67 | 1941.11 | 54.38 |
| **Genotype MS (DF=149)** | 4.17<br>*** | 1.61<br>*** | 7.08<br>*** | 4.80<br>*** | 4.18<br>*** | 3.39<br>*** | 4.76<br>*** | 3.53<br>*** | 3.80<br>*** | 2.83<br>*** | 2.61<br>*** |

*, **, *** and n.s. indicate significance at P ≤ 0.05, 0.01, 0.001 or non-significant effect, respectively.

**Table S2** Ranges of the adjusted ionome traits to variation in GY, P and S under the three environments WL05, WW05 and WW07.

| | GPC | Al | Ca | Cu | Fe | K | Mg | Mn | P | S | Zn |
|---|---|---|---|---|---|---|---|---|---|---|---|
| **WL05** | | | | | | | | | | | |
| **Range GY** | -2.96—2.5 | -1.19—2.48 | -185.55—233.22 | -2.11—3.13 | -16.31—17.45 | -1236.37—1076.27 | -194.66—270.63 | -26.27—41.63 | -685.49—827.87 | -540.5—489.19 | -27.82—36.97 |
| **Range P** | -4.36—3.39 | -1.12—2.66 | -226.42—211.28 | -1.82—3.3 | -17.17—17.29 | -926.68—923.81 | -162.54—202.08 | -27.34—39.55 | --//-- | -474.51—359.05 | -23.53—39.2 |
| **Range S** | -4.83—3.4 | -1.18—2.66 | -181.67—233.5 | -2.13—2.78 | -14.47—15.58 | -1045.22—1380.66 | -190.09—261.1 | -26.78—44.39 | -689.01—957.74 | --//-- | -22.17—33.68 |
| **WW05** | | | | | | | | | | | |
| **Range GY** | -2.33—2.84 | -0.84—1.21 | -211.45—199.92 | -1.95—2.53 | -15.07—32.4 | -928.3—2170.03 | -204.48—209.72 | -20.52—28.98 | -1084.35—834.01 | -352.43—423.74 | -25.27—26.78 |
| **Range P** | -3.9—6.51 | -0.84—1.22 | -200.41—214.63 | -1.58—2.89 | -11.2—35.57 | -1040—1934.62 | -204.3—185.67 | -24.09—33.67 | --//-- | -355.36—431.87 | -19.94—29.67 |
| **Range S** | -2.59—7.04 | -0.89—1.27 | -198.52—208.34 | -2.02—2.53 | -10.9—28.83 | -954.23—2076.15 | -202.55—178.6 | -20.74—29.31 | -1194.78—837.39 | --//-- | -22.27—22.61 |
| **WW07** | | | | | | | | | | | |
| **Range GY** | -2.82—4.01 | -1.64—2.79 | -126—179.26 | -2.32—3.96 | -8.29—12.62 | -664.67—1077.83 | -238.38—245.39 | -12.53—10.21 | -756.92—507.32 | -332.62—366.34 | -16.08—21.53 |
| **Range P** | -3.01—3.61 | -1.51—2.81 | -120.36—186.11 | -12.55—10.65 | -8.36—10.72 | -644.15—1113.39 | -144.26—188.97 | -2.3—3.5 | --//-- | -275.59—243.62 | -16.4—24.81 |
| **Range S** | -2.7—3.64 | -1.51—2.87 | -104.05—192.91 | -13.14—9.45 | -7.87—9.8 | -733.31—1144.16 | -158.49—197.41 | -2.57—3.56 | -508.85—473.4 | --//-- | -19.18—18.84 |

**Table S3** Coefficients of correlation (r) between ionome traits under WL05. Significant correlation coefficients are marked in red (P<0.05).

| | GPC | Al | Ca | Cu | Fe | K | Mg | Mn | P | S | Zn |
|---|---|---|---|---|---|---|---|---|---|---|---|
| GPC | 1.00 | 0.07 | 0.17 | 0.48 | 0.52 | 0.31 | 0.43 | 0.22 | 0.56 | 0.50 | 0.58 |
| Al | 0.07 | 1.00 | 0.33 | 0.23 | 0.25 | 0.26 | 0.18 | -0.04 | 0.19 | 0.03 | 0.20 |
| Ca | 0.17 | 0.33 | 1.00 | 0.05 | 0.14 | 0.53 | 0.54 | 0.28 | 0.46 | 0.28 | 0.22 |
| Cu | 0.48 | 0.23 | 0.05 | 1.00 | 0.65 | 0.22 | 0.29 | 0.15 | 0.40 | 0.42 | 0.60 |
| Fe | 0.52 | 0.25 | 0.14 | 0.65 | 1.00 | 0.26 | 0.28 | 0.09 | 0.41 | 0.41 | 0.76 |
| K | 0.31 | 0.26 | 0.53 | 0.22 | 0.26 | 1.00 | 0.40 | 0.08 | 0.63 | 0.08 | 0.25 |
| Mg | 0.43 | 0.18 | 0.54 | 0.29 | 0.28 | 0.40 | 1.00 | 0.34 | 0.70 | 0.52 | 0.40 |
| Mn | 0.22 | -0.04 | 0.28 | 0.15 | 0.09 | 0.08 | 0.34 | 1.00 | 0.18 | 0.33 | 0.34 |
| P | 0.56 | 0.19 | 0.46 | 0.40 | 0.41 | 0.63 | 0.70 | 0.18 | 1.00 | 0.48 | 0.46 |
| S | 0.50 | 0.03 | 0.28 | 0.42 | 0.41 | 0.08 | 0.52 | 0.33 | 0.48 | 1.00 | 0.59 |
| Zn | 0.58 | 0.20 | 0.22 | 0.60 | 0.76 | 0.25 | 0.40 | 0.34 | 0.46 | 0.59 | 1.00 |

**Table S4** Coefficients of correlation (r) between ionome traits under WW05. Significant correlation coefficients are marked in red (P<0.05).

| | GPC | Al | Ca | Cu | Fe | K | Mg | Mn | P | S | Zn |
|---|---|---|---|---|---|---|---|---|---|---|---|
| GPC | 1.00 | 0.13 | 0.21 | 0.46 | 0.57 | 0.02 | 0.53 | 0.20 | 0.45 | 0.60 | 0.56 |
| Al | 0.13 | 1.00 | 0.31 | 0.12 | 0.15 | 0.24 | 0.19 | 0.04 | 0.14 | 0.16 | 0.12 |
| Ca | 0.21 | 0.31 | 1.00 | 0.07 | 0.12 | 0.19 | 0.39 | 0.12 | 0.36 | 0.23 | 0.17 |
| Cu | 0.46 | 0.12 | 0.07 | 1.00 | 0.51 | 0.19 | 0.38 | 0.12 | 0.31 | 0.29 | 0.44 |
| Fe | 0.57 | 0.15 | 0.12 | 0.51 | 1.00 | -0.01 | 0.43 | 0.12 | 0.44 | 0.50 | 0.79 |
| K | 0.02 | 0.24 | 0.19 | 0.19 | -0.01 | 1.00 | 0.28 | 0.11 | 0.39 | 0.04 | -0.01 |
| Mg | 0.53 | 0.19 | 0.39 | 0.38 | 0.43 | 0.28 | 1.00 | 0.22 | 0.67 | 0.55 | 0.48 |
| Mn | 0.20 | 0.04 | 0.12 | 0.12 | 0.12 | 0.11 | 0.22 | 1.00 | 0.19 | 0.01 | 0.22 |
| P | 0.45 | 0.14 | 0.36 | 0.31 | 0.44 | 0.39 | 0.67 | 0.19 | 1.00 | 0.42 | 0.57 |
| S | 0.60 | 0.16 | 0.23 | 0.29 | 0.50 | 0.04 | 0.55 | 0.01 | 0.42 | 1.00 | 0.52 |
| Zn | 0.56 | 0.12 | 0.17 | 0.44 | 0.79 | -0.01 | 0.48 | 0.22 | 0.57 | 0.52 | 1.00 |

**Table S5** Coefficients of correlation (r) between ionome traits under WW07. Significant correlation coefficients are marked in red ($P<0.05$).

| | GPC | Al | Ca | Cu | Fe | K | Mg | Mn | P | S | Zn |
|---|---|---|---|---|---|---|---|---|---|---|---|
| GPC | 1.00 | 0.11 | 0.02 | 0.36 | 0.36 | -0.13 | 0.63 | 0.32 | 0.50 | 0.66 | 0.47 |
| Al | 0.11 | 1.00 | 0.18 | 0.09 | 0.23 | 0.26 | 0.14 | 0.11 | 0.17 | 0.09 | 0.07 |
| Ca | 0.02 | 0.18 | 1.00 | 0.14 | 0.27 | 0.01 | 0.17 | 0.21 | 0.34 | 0.35 | 0.32 |
| Cu | 0.36 | 0.09 | 0.14 | 1.00 | 0.44 | 0.03 | 0.38 | 0.38 | 0.40 | 0.36 | 0.61 |
| Fe | 0.36 | 0.23 | 0.27 | 0.44 | 1.00 | -0.02 | 0.28 | 0.52 | 0.44 | 0.41 | 0.69 |
| K | -0.13 | 0.26 | 0.01 | 0.03 | -0.02 | 1.00 | -0.03 | -0.11 | 0.17 | -0.07 | -0.18 |
| Mg | 0.63 | 0.14 | 0.17 | 0.38 | 0.28 | -0.03 | 1.00 | 0.20 | 0.65 | 0.64 | 0.36 |
| Mn | 0.32 | 0.11 | 0.21 | 0.38 | 0.52 | -0.11 | 0.20 | 1.00 | 0.18 | 0.29 | 0.49 |
| P | 0.50 | 0.17 | 0.34 | 0.40 | 0.44 | 0.17 | 0.65 | 0.18 | 1.00 | 0.65 | 0.53 |
| S | 0.66 | 0.09 | 0.35 | 0.36 | 0.41 | -0.07 | 0.64 | 0.29 | 0.65 | 1.00 | 0.53 |
| Zn | 0.47 | 0.07 | 0.32 | 0.61 | 0.69 | -0.18 | 0.36 | 0.49 | 0.53 | 0.53 | 1.00 |

**Table S6** Coefficients of correlation (r) between the adjusted ionome traits to variation in GY under WL05 (above diagonal of table)and coefficients of correlation (r) between the adjusted ionome traits to variation in GY under WW05 (below diagonal of table). Significant correlation coefficients are marked red text (P<0.05).

| | | WL05 | | | | | | | | | | |
|---|---|---|---|---|---|---|---|---|---|---|---|---|
| | | GPD | Al | Ca | Cu | Fe | K | Mg | Mn | P | S | Zn |
| | GPD | 1.00 | 0.04 | 0.22 | 0.45 | 0.45 | 0.25 | 0.48 | 0.26 | 0.52 | 0.49 | 0.56 |
| | Al | 0.18 | 1.00 | 0.33 | 0.16 | 0.15 | 0.23 | 0.18 | -0.02 | 0.13 | -0.01 | 0.13 |
| | Ca | 0.18 | 0.30 | 1.00 | -0.03 | 0.11 | 0.53 | 0.57 | 0.25 | 0.49 | 0.26 | 0.21 |
| | Cu | 0.46 | 0.14 | 0.09 | 1.00 | 0.57 | 0.09 | 0.21 | 0.12 | 0.29 | 0.35 | 0.54 |
| | Fe | 0.49 | 0.15 | 0.07 | 0.46 | 1.00 | 0.09 | 0.21 | 0.12 | 0.20 | 0.31 | 0.68 |
| WW05 | K | -0.01 | 0.24 | 0.17 | 0.15 | -0.10 | 1.00 | 0.38 | 0.07 | 0.57 | -0.01 | 0.10 |
| | Mg | 0.50 | 0.13 | 0.28 | 0.32 | 0.29 | 0.19 | 1.00 | 0.35 | 0.72 | 0.49 | 0.38 |
| | Mn | 0.22 | 0.05 | 0.09 | 0.19 | 0.16 | 0.13 | 0.27 | 1.00 | 0.24 | 0.37 | 0.42 |
| | P | 0.33 | 0.09 | 0.29 | 0.25 | 0.32 | 0.36 | 0.60 | 0.17 | 1.00 | 0.41 | 0.30 |
| | S | 0.60 | 0.13 | 0.18 | 0.23 | 0.45 | -0.04 | 0.49 | 0.04 | 0.31 | 1.00 | 0.57 |
| | Zn | 0.50 | 0.07 | 0.09 | 0.41 | 0.75 | -0.06 | 0.37 | 0.26 | 0.49 | 0.49 | 1.00 |

**Table S7** Coefficients of correlation (r) between the adjusted ionome traits to variation in GY under WW07. Significant correlation coefficients are marked red text (P<0.05).

| | | WW07 | | | | | | | | | | |
|---|---|---|---|---|---|---|---|---|---|---|---|---|
| | | GPD | Al | Ca | Cu | Fe | K | Mg | Mn | P | S | Zn |
| WW07 | GPD | 1.00 | 0.08 | 0.06 | 0.38 | 0.37 | -0.22 | 0.61 | 0.42 | 0.48 | 0.65 | 0.52 |
| | Al | 0.08 | 1.00 | 0.20 | 0.09 | 0.23 | 0.23 | 0.11 | 0.16 | 0.15 | 0.07 | 0.09 |
| | Ca | 0.06 | 0.20 | 1.00 | 0.14 | 0.28 | 0.05 | 0.20 | 0.19 | 0.37 | 0.38 | 0.31 |
| | Cu | 0.38 | 0.09 | 0.14 | 1.00 | 0.44 | 0.04 | 0.40 | 0.39 | 0.41 | 0.37 | 0.61 |
| | Fe | 0.37 | 0.23 | 0.28 | 0.44 | 1.00 | -0.02 | 0.28 | 0.55 | 0.44 | 0.42 | 0.70 |
| | K | -0.22 | 0.23 | 0.05 | 0.04 | -0.02 | 1.00 | -0.11 | -0.04 | 0.13 | -0.13 | -0.15 |
| | Mg | 0.61 | 0.11 | 0.20 | 0.40 | 0.28 | -0.11 | 1.00 | 0.29 | 0.63 | 0.63 | 0.41 |
| | Mn | 0.42 | 0.16 | 0.19 | 0.39 | 0.55 | -0.04 | 0.29 | 1.00 | 0.25 | 0.36 | 0.48 |
| | P | 0.48 | 0.15 | 0.37 | 0.41 | 0.44 | 0.13 | 0.63 | 0.25 | 1.00 | 0.64 | 0.56 |
| | S | 0.65 | 0.07 | 0.38 | 0.37 | 0.42 | -0.13 | 0.63 | 0.36 | 0.64 | 1.00 | 0.56 |
| | Zn | 0.52 | 0.09 | 0.31 | 0.61 | 0.70 | -0.15 | 0.41 | 0.48 | 0.56 | 0.56 | 1.00 |

**Table S8** Coefficients of correlation (r) between the adjusted ionome traits to variation in P (below diagonal of table) and coefficients of correlation (r) between the adjusted ionome traits to variation in S (above diagonal of table) under WL05. Significant correlation coefficients are marked in red (P<0.05).

| | | S | | | | | | | | | |
|---|---|---|---|---|---|---|---|---|---|---|---|
| | | GPC | Al | Ca | Cu | Fe | K | Mg | Mn | P | Zn |
| P | GPC | 1.00 | 0.07 | 0.04 | 0.35 | 0.40 | 0.31 | 0.22 | 0.06 | 0.42 | 0.40 |
| | Al | -0.04 | 1.00 | 0.34 | 0.24 | 0.26 | 0.25 | 0.19 | -0.05 | 0.20 | 0.23 |
| | Ca | -0.11 | 0.28 | 1.00 | -0.08 | 0.04 | 0.53 | 0.48 | 0.21 | 0.39 | 0.07 |
| | Cu | 0.34 | 0.17 | -0.17 | 1.00 | 0.58 | 0.21 | 0.09 | 0.01 | 0.25 | 0.48 |
| | Fe | 0.38 | 0.20 | -0.06 | 0.58 | 1.00 | 0.25 | 0.08 | -0.06 | 0.27 | 0.71 |
| | K | -0.07 | 0.18 | 0.34 | -0.05 | 0.00 | 1.00 | 0.42 | 0.06 | 0.68 | 0.25 |
| | Mg | 0.06 | 0.07 | 0.34 | 0.02 | -0.01 | -0.08 | 1.00 | 0.21 | 0.60 | 0.14 |
| | Mn | 0.14 | -0.08 | 0.23 | 0.09 | 0.01 | -0.05 | 0.30 | 1.00 | 0.03 | 0.20 |
| | S | 0.32 | -0.07 | 0.07 | 0.29 | 0.27 | -0.33 | 0.30 | 0.28 | 1.00 | 0.25 |
| | Zn | 0.44 | 0.13 | 0.01 | 0.51 | 0.71 | -0.06 | 0.13 | 0.30 | 0.48 | 1.00 |

**Table S9** Coefficients of correlation (r) between the adjusted ionome traits to variation in P (below diagonal of table) and coefficients of correlation (r) between the adjusted ionome traits to variation in S (above diagonal of table) under WW05. Significant correlation coefficients are marked in red (P<0.05).

| | | S | | | | | | | | | |
|---|---|---|---|---|---|---|---|---|---|---|---|
| | | GPC | Al | Ca | Cu | Fe | K | Mg | Mn | S | Zn |
| P | GPC | 1.00 | 0.04 | 0.10 | 0.37 | 0.40 | 0.00 | 0.31 | 0.24 | 0.27 | 0.37 |
| | Al | 0.07 | 1.00 | 0.29 | 0.07 | 0.09 | 0.24 | 0.13 | 0.04 | 0.08 | 0.05 |
| | Ca | 0.06 | 0.28 | 1.00 | 0.00 | 0.00 | 0.19 | 0.32 | 0.12 | 0.30 | 0.06 |
| | Cu | 0.38 | 0.08 | -0.05 | 1.00 | 0.44 | 0.19 | 0.27 | 0.12 | 0.22 | 0.35 |
| | Fe | 0.47 | 0.10 | -0.05 | 0.44 | 1.00 | -0.04 | 0.21 | 0.14 | 0.30 | 0.72 |
| | K | -0.18 | 0.20 | 0.06 | 0.08 | -0.22 | 1.00 | 0.31 | 0.11 | 0.41 | -0.04 |
| | Mg | 0.35 | 0.13 | 0.21 | 0.24 | 0.19 | 0.04 | 1.00 | 0.26 | 0.58 | 0.27 |
| | Mn | 0.13 | 0.02 | 0.05 | 0.06 | 0.05 | 0.04 | 0.13 | 1.00 | 0.20 | 0.25 |
| | S | 0.51 | 0.11 | 0.10 | 0.19 | 0.39 | -0.15 | 0.40 | -0.08 | 1.00 | 0.45 |
| | Zn | 0.42 | 0.05 | -0.04 | 0.34 | 0.73 | -0.30 | 0.16 | 0.14 | 0.38 | 1.00 |

**Table S10** Coefficients of correlation (r) between the adjusted ionome traits to variation in P (below diagonal of table) and coefficients of correlation (r) between the adjusted ionome traits to variation in S (above diagonal of table) under WW07. Significant correlation coefficients are marked in red (P<0.05).

| | | S | | | | | | | | | |
|---|---|---|---|---|---|---|---|---|---|---|---|
| | | GPC | Al | Ca | Cu | Fe | K | Mg | Mn | P | Zn |
| P | GPC | 1.00 | 0.07 | -0.30 | 0.18 | 0.13 | -0.10 | 0.36 | 0.18 | 0.13 | 0.19 |
| | Al | 0.03 | 1.00 | 0.16 | 0.06 | 0.21 | 0.26 | 0.11 | 0.09 | 0.14 | 0.03 |
| | Ca | -0.19 | 0.13 | 1.00 | 0.02 | 0.15 | 0.04 | -0.09 | 0.12 | 0.16 | 0.17 |
| | Cu | 0.20 | 0.03 | 0.01 | 1.00 | 0.35 | 0.06 | 0.21 | 0.30 | 0.24 | 0.53 |
| | Fe | 0.18 | 0.17 | 0.15 | 0.32 | 1.00 | 0.02 | 0.02 | 0.46 | 0.24 | 0.61 |
| | K | -0.25 | 0.23 | -0.05 | -0.04 | -0.10 | 1.00 | 0.02 | -0.09 | 0.29 | -0.17 |
| | Mg | 0.46 | 0.04 | -0.08 | 0.17 | -0.01 | -0.19 | 1.00 | 0.02 | 0.40 | 0.04 |
| | Mn | 0.27 | 0.08 | 0.16 | 0.34 | 0.50 | -0.15 | 0.11 | 1.00 | -0.01 | 0.41 |
| | S | 0.50 | -0.02 | 0.18 | 0.14 | 0.19 | -0.25 | 0.38 | 0.23 | 1.00 | 0.29 |
| | Zn | 0.28 | -0.02 | 0.18 | 0.51 | 0.60 | -0.32 | 0.03 | 0.47 | 0.28 | 1.00 |

**Table S11** Kendall's Tau coefficients of rank correlations between the initial and the adjusted traits under the three environments WL05, WW05 and WW07.

| Initial traits | Environments | Adjusted traits | | |
|---|---|---|---|---|
| | | GY | P | S |
| GPC | WL05 | 0.791 | 0.791 | 0.848 |
| | WW05 | 0.669 | 0.873 | 0.789 |
| | WW07 | 0.851 | 0.857 | 0.713 |
| Al | WL05 | 0.833 | 0.974 | 0.999 |
| | WW05 | 0.992 | 0.985 | 0.982 |
| | WW07 | 0.910 | 0.977 | 0.990 |
| Ca | WL05 | 0.954 | 0.885 | 0.958 |
| | WW05 | 0.869 | 0.923 | 0.966 |
| | WW07 | 0.932 | 0.918 | 0.927 |
| Cu | WL05 | 0.805 | 0.908 | 0.908 |
| | WW05 | 0.856 | 0.949 | 0.951 |
| | WW07 | 0.990 | 0.880 | 0.908 |
| Fe | WL05 | 0.667 | 0.898 | 0.912 |
| | WW05 | 0.689 | 0.826 | 0.831 |
| | WW07 | 0.991 | 0.880 | 0.893 |
| K | WL05 | 0.771 | 0.774 | 0.996 |
| | WW05 | 0.924 | 0.903 | 0.999 |
| | WW07 | 0.821 | 0.978 | 0.996 |
| Mg | WL05 | 0.974 | 0.668 | 0.833 |
| | WW05 | 0.717 | 0.744 | 0.832 |
| | WW07 | 0.856 | 0.756 | 0.763 |
| Mn | WL05 | 0.925 | 0.987 | 0.954 |
| | WW05 | 0.987 | 0.971 | 1.000 |
| | WW07 | 0.845 | 0.982 | 0.949 |
| P | WL05 | 0.767 | --//-- | 0.861 |
| | WW05 | 0.711 | --//-- | 0.876 |
| | WW07 | 0.900 | --//-- | 0.767 |
| S | WL05 | 0.894 | 0.857 | --//-- |
| | WW05 | 0.815 | 0.884 | --//-- |
| | WW07 | 0.913 | 0.763 | --//-- |
| Zn | WL05 | 0.745 | 0.880 | 0.802 |
| | WW05 | 0.731 | 0.774 | 0.839 |
| | WW07 | 0.926 | 0.823 | 0.822 |
